# Unfolded protein response is a critical mediator in *Campylobacter jejuni* pathogenesis and host defence

**DOI:** 10.1101/2025.01.31.635839

**Authors:** Geunhye Hong, Zahra Omole, Cadi Davies, Janie Liaw, Anna D. Grabowska, Barbara Canonico, Nicolae Corcionivoschi, Brendan W. Wren, Ezra Aksoy, Nick Dorrell, Abdi Elmi, Ozan Gundogdu

## Abstract

*Campylobacter jejuni* is the major bacterial cause of foodborne gastroenteritis worldwide. How this pathogen interacts with the host defence machinery of human intestinal epithelial cells (IECs) and is involved in pathogenesis remains elusive. Bacterial pathogens utilise strategies to gain access to the eukaryotic cell machinery that can involve subversion of biological processes in host. Unfolded protein response (UPR) is a highly conserved host cell stress response to the accumulation of misfolded proteins in the endoplasmic reticulum (ER) and is a conserved evolutionary response against invading pathogens. Several bacterial pathogens can induce the UPR for their own survival and thus design a dual scenario where UPR can both protect and facilitate pathogen evasion.

Herein, we investigated whether UPR represents a virulence mechanism exploited by *C. jejuni* during bacterial invasion in human IECs. Our data show that following *C. jejuni* infection, we observe upregulation of protein kinase R-like ER kinase (PERK), inositol-requiring enzyme 1α (IRE1α) and activating transcription factor 6 (ATF6) pathways of the UPR, albeit differentially in human T84 and Caco-2 cells. Chemical induction of UPR by thapsigargin in host cells reduced intracellular survival of *C. jejuni* while conversely pretreatment with UPR inhibitors increased intracellular survival of *C. jejuni* and attenuated IL-8 release. Finally, we show that *C. jejuni* mutant *C. jejuni* capsular polysaccharide and flagella contribute to UPR activation in IECs. Collectively, these findings provide mechanistic insights into how *C. jejuni* infection leads to UPR activation and inflammation, potentially contributing to downstream *C. jejuni*-mediated damage.

**Author Summary:** *Campylobacter jejuni* is one of the most prevalent bacterial causes of foodborne human gastroenteritis worldwide. The common symptoms of campylobacteriosis include diarrhoea, abdominal pain and fever. *C. jejuni*-induced diarrhoea is associated with disruption of tight junctions and adherens junctions, downregulation of epithelial sodium channels and disruption of bile reabsorption. In addition, *C. jejuni*-induced diarrhoea in part result from intestinal inflammation caused by increased *C. jejuni* colonisation and invasion due to dysfunctional intestinal barrier. *C. jejuni* was reported to cause an inflammatory signalling cascade via ligand recognition by Toll-like receptor 2 (TLR2), TLR4 and nucleotide-binding oligomerisation domain (NOD) in IECs. However, other potential mechanisms of *C. jejuni*-induced inflammation, with respect to the IEC responses remains unknown. In this study, we demonstrate UPR activation following *C. jejuni* infection. The UPR is a conserved eukaryotic regulatory response which is initiated to relieve ER stress and recover ER homeostasis, however several bacteria have evolved to exploit UPR pathways for their own survival and replication, creating a dual scenario where UPR can both protect and facilitate pathogen evasion. This study shows that the activation of the PERK pathway of host UPR is activated in the presence of live *C. jejuni* or its structural components, flagella and a capsular polysaccharide. Furthermore, we propose that although activation of the UPR is a conserved evolutionary response against invading pathogens, UPR induction induces inflammation which potentially contributes to *C. jejuni*-mediated damage.

## Introduction

All living cells require an optimal environment to maintain normal cellular function and metabolism [1]. The endoplasmic reticulum (ER) is the largest eukaryotic organelle which stores Ca^2+^ and processes lipid and protein synthesis, protein folding and transport [2, 3]. Disruption of cellular protein folding increases the amount of misfolded proteins within the ER lumen, known as ER stress [4]. As the ER plays a significant role in cellular physiology, ER homeostasis is tightly regulated by a coordinated system [4, 5]. The unfolded protein response (UPR) is an eukaryotic stress surveillance system which is activated to relieve ER stress and recover ER homeostasis [4].

ER membrane contains three transmembrane proteins, protein kinase R-like ER kinase (PERK), inositol-requiring enzyme 1α (IRE1α) and activating transcription factor 6 (ATF6) which detect the unfolded proteins within the ER [4, 5]. The activity of the UPR sensors is controlled by binding and detachment of the binding-immunoglobulin protein (BiP). Under stabilised conditions, BiP is bound to the luminal domain of the UPR sensors to stabilise their inactive state. Whereas, during ER stress, BiP is detached from the UPR sensors due to its higher affinity to unfolded proteins, which leads to conformational changes of the UPR sensors leading to UPR activation [4, 5]. UPR activation induces expression of proteins associated with ER chaperons, ER-associated degradation (ERAD), amino acid transport and metabolism and resistance to oxidative stress [4, 5]. In addition to restoration of cellular physiology within the ER, UPR activation is closely interconnected with proinflammatory responses against bacterial infection [6].

*Campylobacter jejuni* is the major *Campylobacter* species that causes human gastroenteritis worldwide [7]. The predominant symptoms include watery or bloody diarrhoea, vomiting, abdominal pain, and fever [8, 9]. *C. jejuni* infection is also implicated in development of Guillain-Barré Syndrome (GBS) and Miller-Fisher syndrome that rare but severe post-infectious autoimmune diseases of the peripheral nervous system [10]. Notably, *C. jejuni* infection can lead to developmental impairment and death of children in low- and middle-income countries. [11]. Despite of its prevalence and importance, our understanding of how *C. jejuni* causes diarrhoeal disease is fragmented and incomplete. We know of four broad mechanisms that can lead to diarrhoea upon *C. jejuni* infection of humans; (1) downregulation of epithelial sodium (Na^+^) channel expression [12]; (2) prevention of reabsorption of bile acids[13]; (3) disruption of intestinal barrier by cleaving tight junctions and adherens junction proteins [14–16]; and (4) induction of intestinal inflammation [17–19]. *C. jejuni* is known to induce inflammation via pattern recognition receptor (PRR) detection such as Toll-like receptor 2 (TLR2), Toll-like receptor 4 (TLR4) and nucleotide-binding oligomerisation domain 1 (NOD1) [17–19]. Yet the molecular and cellular mechanisms of how *C. jejuni* infection promotes inflammation remain unclear.

UPR activation enhances host defence mechanisms against bacterial pathogens through the activation of proinflammatory responses [20]. Whereas some pathogens such as *Salmonella enterica*, *Brucella melitensis* and *Chlamydia* species have evolved to exploit UPR pathways for their own survival and replication creating a dual scenario where UPR can both protect the host and facilitate pathogen evasion [21–23]. These differential effects of the UPR on bacterial pathogens depend on the unique pathogenic strategies that each bacterial species possesses. Notably, *C. jejuni* has recently been reported to induce the activation of the PERK pathway in human IECs [24]. However, the mechanisms of *C. jejuni*-mediated UPR activation and the significance of UPR activation on *C. jejuni* pathogenesis and *C. jejuni*-mediated inflammation are still to be determined.

In this study, to address the gaps in our knowledge, we conducted a thorough investigation of UPR activation by *C. jejuni* in human IECs. We examined whether *C. jejuni* strains activate all three branches of the UPR (PERK, IRE1α, and ATF6) in T84 and Caco-2 cells. Our findings reveal that all *C. jejuni* strains consistently activated both the PERK and IRE1α pathways in T84 and Caco-2 cells. However, activation of the ATF6 pathway showed strain- and cell line-dependent variations, suggesting a more complex regulatory mechanism for this branch of the UPR. We additionally investigated the contribution of specific *C. jejuni* components on the PERK and IRE1α pathways. We confirmed the role of *C. jejuni* capsule and flagella in the activation of the PERK pathway, corroborating the contribution of the bacterial components in a sole branch of the UPR pathway, yet broader activation of the UPR by live *C. jejuni* which occurred independently of Ca^2+^ release from the ER. Importantly our data showed that UPR activation serves as a host defense mechanism against *C. jejuni* invasion, effectively limiting bacterial growth within IECs. Intriguingly, we also found that UPR activation is simultaneously associated with *C. jejuni*-induced inflammation in human IECs, revealing a dual role for this stress response.

## Results

### *Campylobacter jejuni* activates all three branches of the UPR in T84 and Caco-2 cells

To investigate inter-strain variation in *C. jejuni*-activated UPR, T84 and Caco-2 cells were infected with *C. jejuni* 11168H, 81-176 or 488 strains and mRNA levels of UPR-related genes were measured (Figs 1 and 2). *C. jejuni* 11168H, 81-176 and 488 strains significantly increased mRNA levels of *CHOP* and spliced *XBP1* in T84 cells at 6- and 24-h post-infection (Figs 1A and 1B), indicating PERK and IRE1α pathway activation respectively. A similar pattern was observed for *CHOP* and spliced *XBP1* in Caco-2 cells infected with *C. jejuni* (Figs 2A and 2B). A distinct pattern was observed for *ATF6* in T84 cells at 6 h post-infection with 81-176 (Fig 1C). Unlike 11168H and 488, 81-176 upregulated *ATF6* in T84 cells. In contrast, 81-176 did not increase mRNA level of *ATF6* in Caco-2 cells indicating ATF6 activation exhibits strain- and cell line-specific pattern (Fig 2C). All three *C. jejuni* strains downregulated *BiP* at 6 h post-infection and mRNA level of *BiP* was recovered at 24 h post-infection in both T84 and Caco-2 cells (Figs 1D and 2D). Increase in protein levels of CHOP and spliced XBP1 was also demonstrated in both cell lines infected with 11168H, 81-176 or 488 at 24 h post-infection (Fig 3).

**Fig 1.**
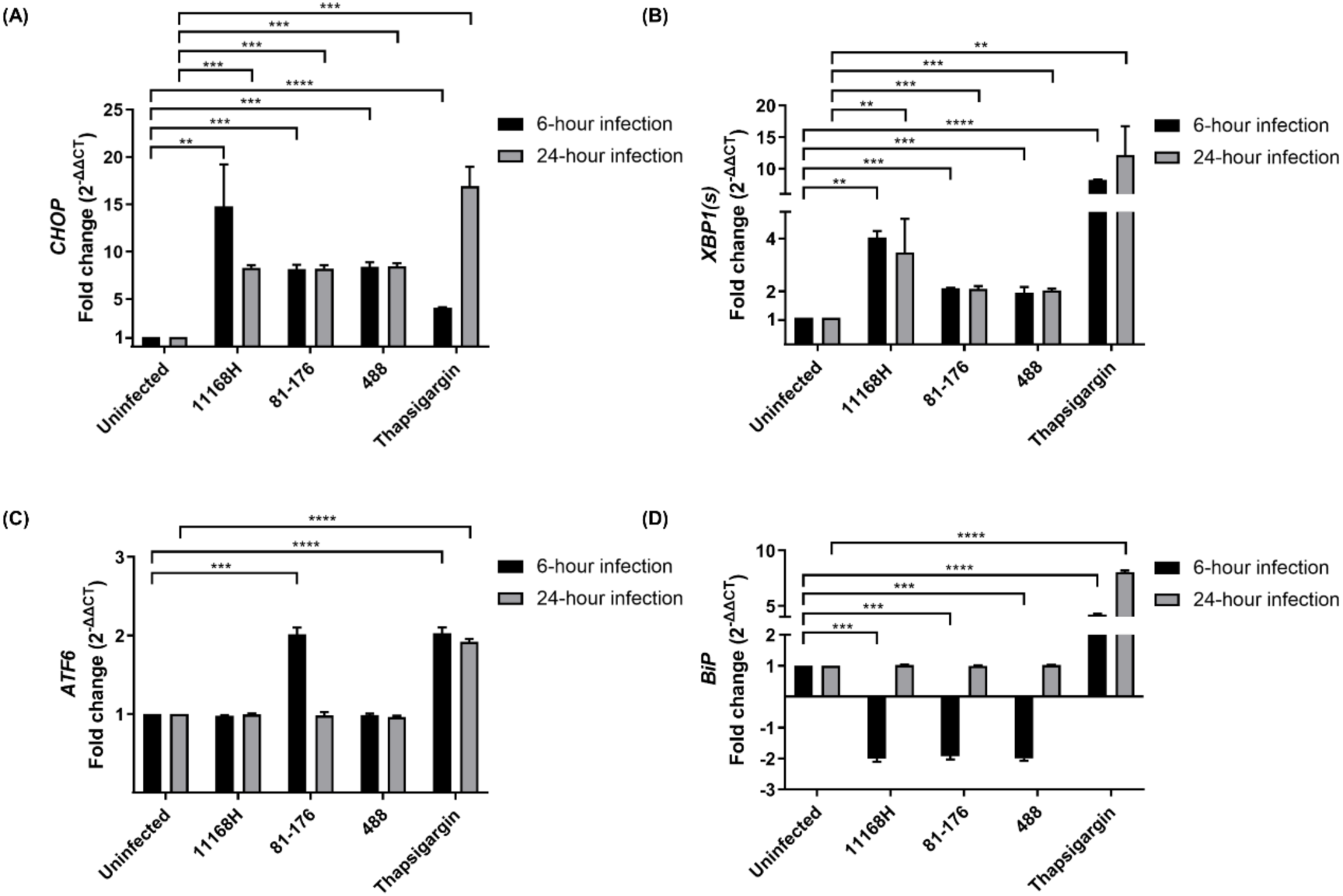
UPR-related gene expression in T84 cells infected with *C. jejuni*. qRT-PCR showing expression of human (A) *CHOP*, (B) spliced *XBP1* [*XBP1(s)*], (C) *ATF6* and (D) *BiP* in T84 cells infected with *C. jejuni* 11168H, 81-176 or 488 wild-type strains for 6 or 24 h at 37°C in a 5% CO_2_ incubator (MOI of 200:1). T84 cells were treated with thapsigargin as a positive control. *GAPDH* was used as an internal control. One sample *t*-test was performed. Asterisks denote a statistically significant difference (** = *p* < 0.01; *** = *p* < 0.001; **** = *p* < 0.0001).

**Fig 2.**
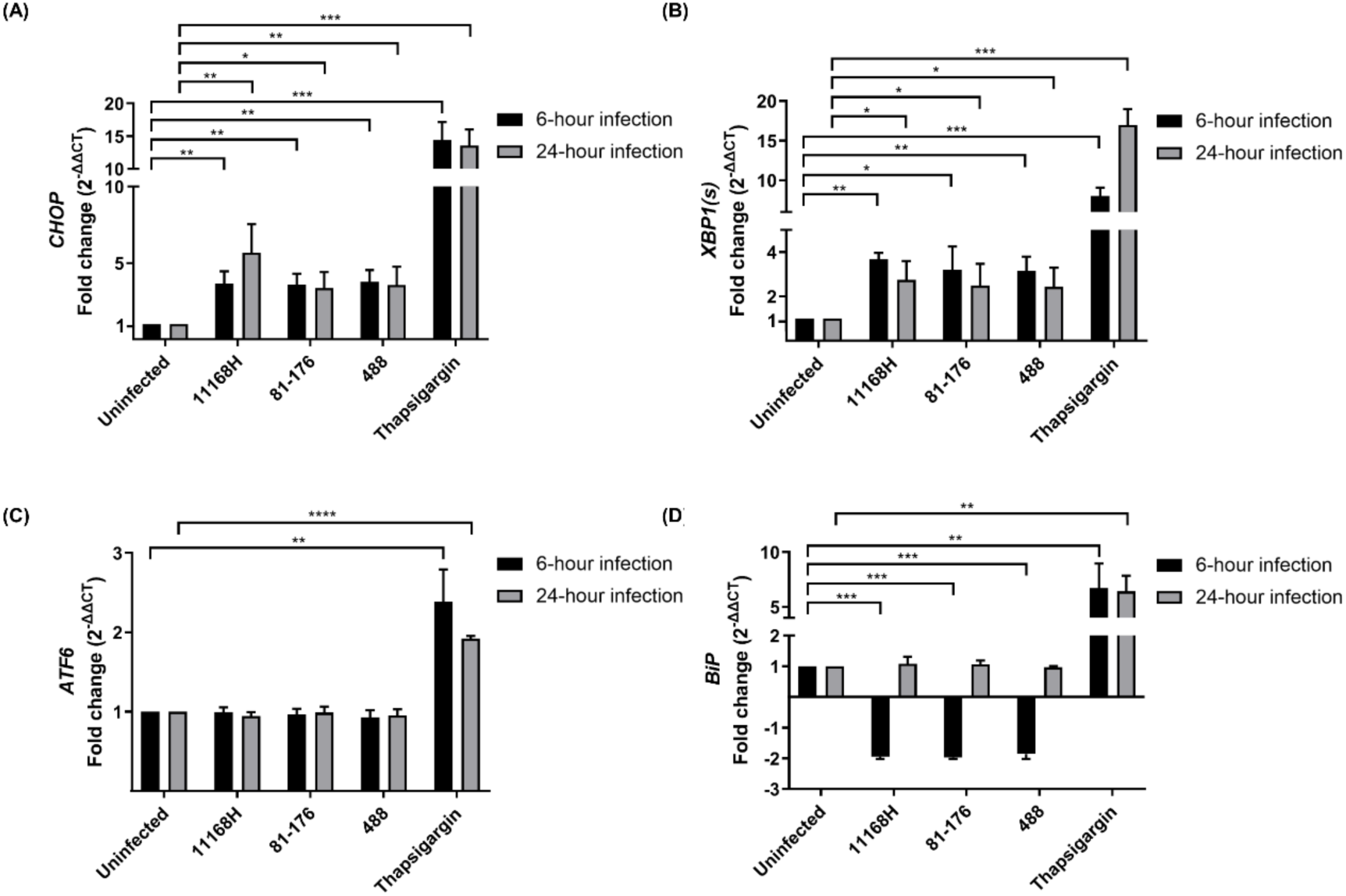
UPR-related gene expression in Caco-2 cells infected with *C. jejuni*. qRT-PCR showing expression of human (A) *CHOP*, (B) spliced *XBP1* [*XBP1(s)*], (C) *ATF6* and (D) *BiP* in Caco-2 cells infected with *C. jejuni* 11168H, 81-176 or 488 wild-type strains for 6 or 24 h at 37°C in a 5% CO_2_ incubator (MOI of 200:1). Caco-2 cells were treated with thapsigargin as a positive control. *GAPDH* was used as an internal control. Three biological and three technical replicates were performed for each experiment. One sample *t*-test was performed. Asterisks denote a statistically significant difference (* = *p* < 0.05; ** = *p* < 0.01; *** = *p* < 0.001; **** = *p* < 0.0001).

**Fig 3.**
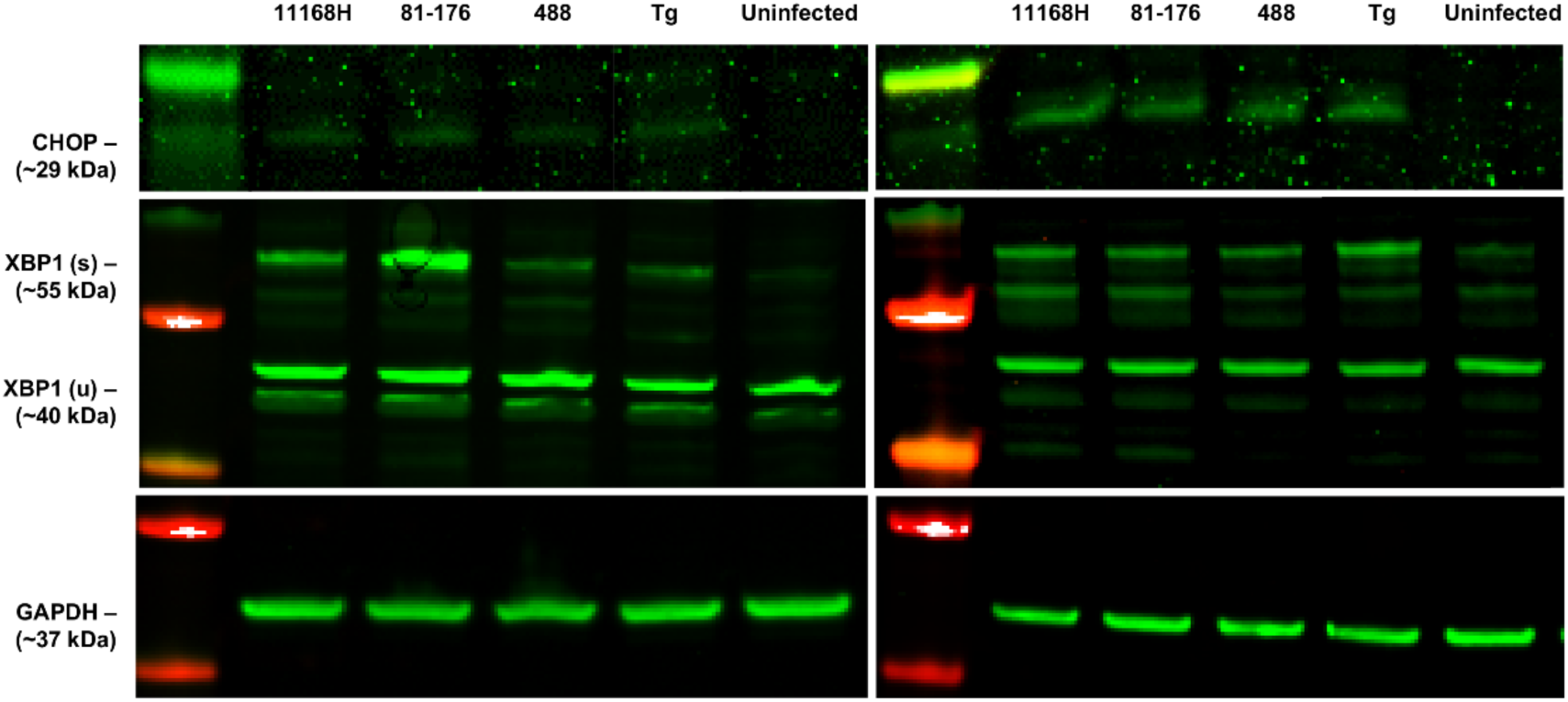
UPR-related protein expression in T84 and Caco-2 cells infected with *C. jejuni*. Western blotting showing protein levels of human CHOP and both spliced & unspliced forms of XBP1 [XBP1(u) & XBP1(s)] in T84 (left panels) and Caco-2 cells (right panels) infected with *C. jejuni* 11168H, 81-176 or 488 wild-type strains for 24 h at 37°C in a 5% CO_2_ incubator (MOI of 200:1). T84 cells were treated with thapsigargin (Tg) as a positive control. GAPDH was used as an internal control.

### The UPR activation is detrimental to *C. jejuni* interaction, invasion and intracellular survival in T84 and Caco-2 cells

The effect of thapsigargin-mediated UPR activation on *C. jejuni* interaction, invasion and intracellular survival in human IECs was explored. Pre-treatment with thapsigargin did not affect *C. jejuni* interaction with or invasion of T84 or Caco-2 cells (Fig 4A-D). In contrast, intracellular survival of *C. jejuni* in both T84 and Caco-2 cells was significantly reduced in thapsigargin-treated cells compared to untreated human IECs (Fig 4E and 4F). To further confirm the impact of the UPR on intracellular *C. jejuni*, using UPR inhibitors, the effect of the UPR on the number of intracellular *C. jejuni* in human IECs was investigated (Fig 5). UPR inhibitors KIRA6, STF-083010 and GSK2656157 significantly downregulated *C. jejuni*-mediated activation PERK and IRE1α pathways in T84 and Caco-2 cells (S1 Fig and S2 Fig). Interestingly, pre-treatment with KIRA6 and STF-083010 significantly increased the number of intracellular *C. jejuni* in T84 and Caco-2 cells (Fig 5A and 5B). Similarly, pre-treatment of GSK2656157 increased the number of intracellular *C. jejuni* in both T84 and Caco-2 cells (Fig 5A and 5B). *C. jejuni* viability with thapsigargin and UPR inhibitors was assessed and each treatment did not affect the viability of *C. jejuni* (S3 Fig). Cytotoxicity of chemical treatments on T84 and Caco-2 cells was also investigated (S4 Fig). Treatment of thapsigargin resulted in 5.7% and 5.5% cytotoxicity on T84 and Caco-2 cells respectively while other treatments did not exert significant cytotoxicity on human IECs (S4 Fig). Collectively, these observations indicate activation of both PERK and IRE1α pathways exhibit a detrimental effect on intracellular *C. jejuni* survival in human IECs.

**Fig 4.**
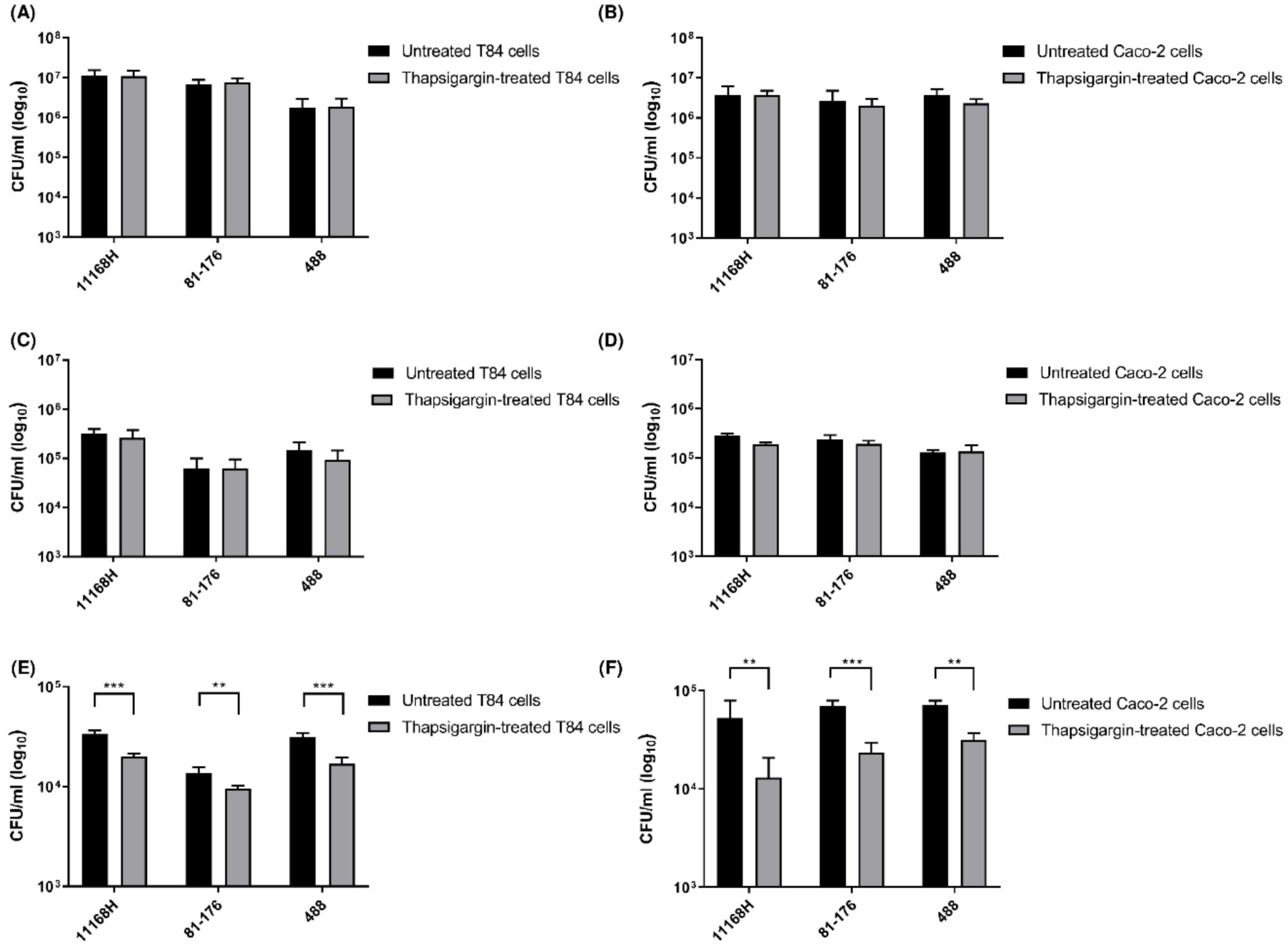
The impact of thapsigargin-induced UPR on C. jejuni interaction, invasion and intracellular survival. T84 or Caco-2 cells were pre-treated with 2 µM of thapsigargin for 6 h and infected with *C. jejuni* 11168H, 81-176 or 488 wild-type strains for 3 h at 37°C in a 5% CO_2_ atmosphere (MOI of 200:1). For interaction assays, (A) T84 or (B) Caco-2 cells were washed with PBS, lysed with 0.1% (v/v) Triton X-100 and CFU/ml were recorded after incubation. For invasion assays, after infection with *C. jejuni*, (C) T84 or (D) Caco-2 cells were incubated with gentamicin (150 µg/ml) for 2 h to kill extracellular bacteria, then lysed with 0.1% (v/v) Triton X-100 and CFU/ml were recorded after incubation. For intracellular survival assays, the 2 h gentamicin treatment as for invasion assays was followed by further incubation with gentamicin (10 µg/ml) for 18 h, then (E) T84 or (F) Caco-2 cells were lysed with 0.1% (v/v) Triton X-100 and CFU/ml were recorded after incubation. One sample *t*-test was performed. Asterisks denote a statistically significant difference (** = *p* < 0.01; *** = *p* < 0.001).

**Fig 5.**
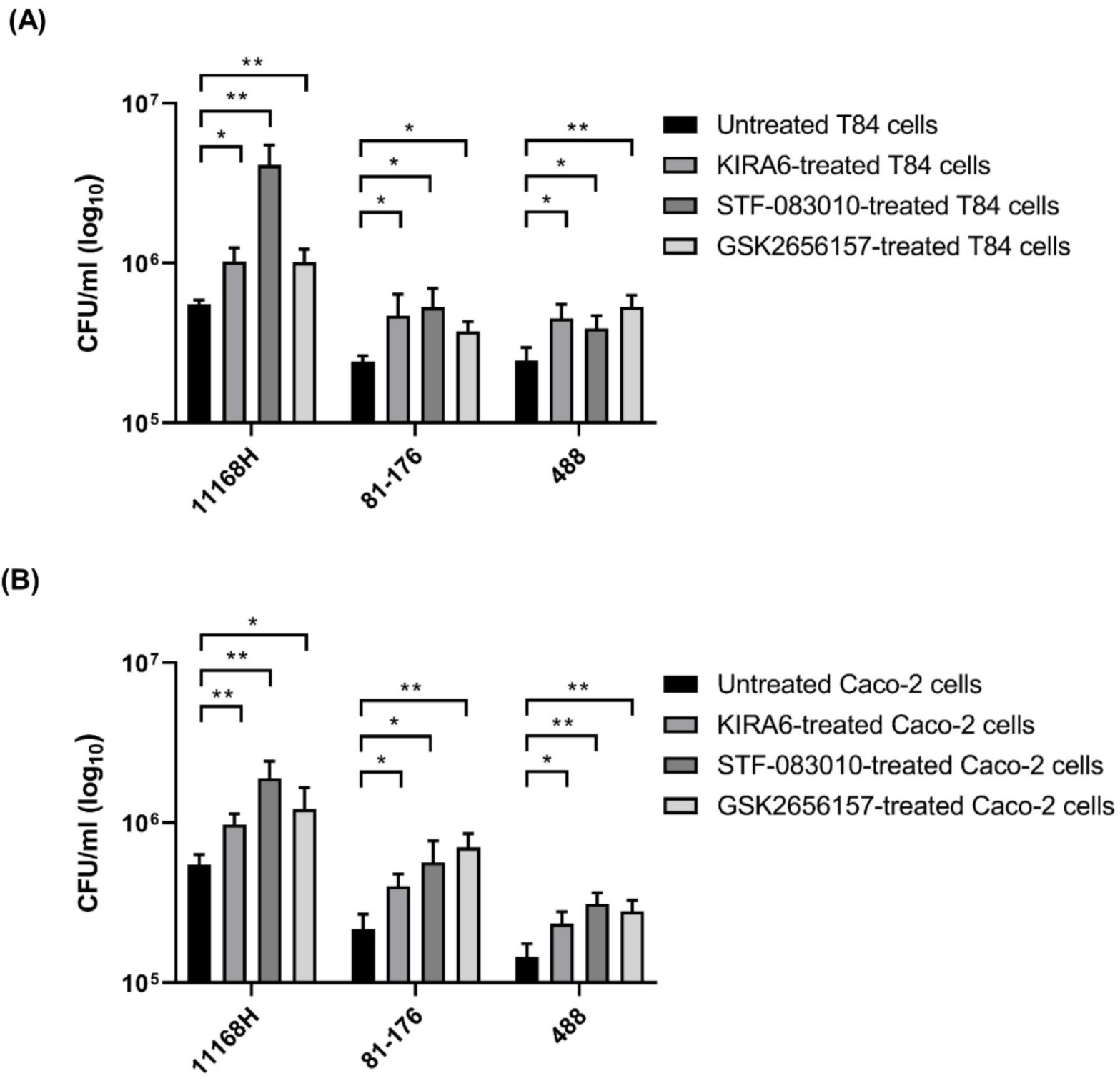
The effect of UPR inhibitors on the number of intracellular *C. jejuni* in human intestinal epithelial cells. (A) T84 or (B) Caco-2 cells were pre-treated with 3 µM of KIRA6, 100 µM of STF-083010 or 3 µM of GSK2656157 for 4 h at 37°C in a 5% CO_2_ incubator. T84 and Caco-2 cells were then infected with *C. jejuni* 11168H, 81-176 or 488 wild-type strains for 24 h at 37°C in a 5% CO_2_ incubator (MOI of 200:1). Human IECs were further treated with 3 µM of KIRA6 and 3 µM of GSK2656157 during *C. jejuni* infection. T84 and Caco-2 cells were washed with PBS and incubated with gentamicin (150 µg/ml) for 2 h to kill extracellular bacteria and then lysed with 0.1% (v/v) Triton X-100 and CFU/ml were recorded after incubation. One sample *t*-test was performed. Asterisks denote a statistically significant difference (* = *p* < 0.05; ** = *p* < 0.01).

### UPR activation contributes to *C. jejuni*-induced inflammation in T84 cells

Given the observed anti-bacterial effects of UPR activation on intracellular *C. jejuni* in T84 and Caco-2 cells and the fact that the UPR is closely linked to inflammatory responses [6], we next explored the relationship between the UPR and *C. jejuni*-induced inflammation. *C. jejuni* 11168H, 81-176 and 488 strains all induced IL-8 secretion in T84 cells at 24 h post-infection (Fig 6A), whereas only 81-176 induced IL-8 secretion at 6 h post-infection suggesting inter-strain variation in IL-8 induction (Fig 6A). Thapsigargin induced IL-8 release from T84 cells at both 6- and 24-h post-treatment indicating thapsigargin-mediated UPR activation is associated with inflammation (Fig 6A). Next, to investigate if *C. jejuni*-induced inflammation is implicated in *C. jejuni*-mediated UPR activation, we assessed the impact of the UPR inhibitors on IL-8 release from T84 cells by *C. jejuni*. Pre-treatment with KIRA6 and GSK2656157 resulted in significantly impaired *C. jejuni*- and thapsigargin-mediated IL-8 secretion (Fig 6B-E). In contrast, pre-treatment with STF-083010 significantly augmented *C. jejuni*-induced IL-8 secretion whereas the treatment did not affect thapsigargin-mediated IL-8 release from T84 cells (Fig 6B-E). These results suggested that IRE1α inhibitors exert differential effects on IL-8 secretion. Overall, we concluded that *C. jejuni*-mediated inflammation in human IECs activate PERK and IRE1α pathways of UPR, yet with opposing effects on host inflammatory response.

**Fig 6.**
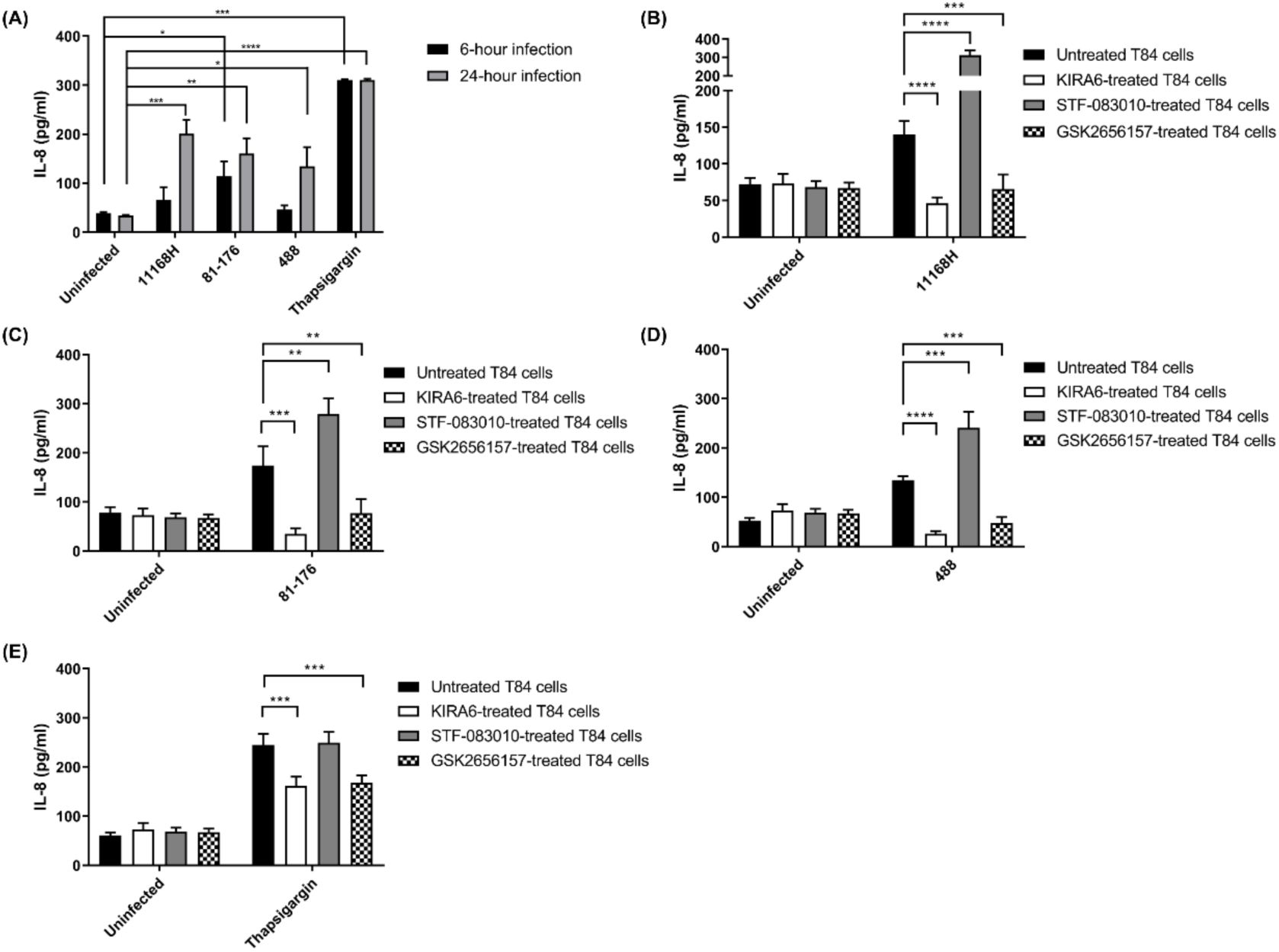
IL-8 release from T84 cells infected with *C. jejuni* or treated with thapsigargin and the impact of UPR inhibitors on IL-8 release from T84 cells infected with *C. jejuni* or treated with thapsigargin. (A) T84 cells in a 24-well plate were infected with *C. jejuni* 11168H, 81-176 or 488 wild-type strains (MOI 200:1) or treated with 2 µM of thapsigargin for 6 or 24 h at 37°C in a 5% CO_2_ incubator. T84 cells were pre-treated with 3 µM of KIRA6, 100 µM of STF-083010 or 3 µM of GSK2656157 for 4 h at 37°C in a 5% CO_2_ incubator. T84 cells were then infected with *C. jejuni* 11168H, 81-176 or 488 wild-type strains (MOI 200:1) or treated with 2 µM of thapsigargin for 24 h at 37°C in a 5% CO_2_ incubator. T84 cells were further treated with 3 µM of KIRA6 and 3 µM of GSK2656157 during *C. jejuni* infection. T84 cells in a 24-well plate were infected with *C. jejuni* (B) 11168H, (C) 81-176 or (D) 488 wild-type strains or treated with (E) thapsigargin for 6 or 24 h at 37°C in a 5% CO_2_ incubator. Medium from each well was subjected to human IL-8 ELISA to measure the concentrations of IL-8. One sample *t*-test was performed. Asterisks denote a statistically significant difference (* = *p* < 0.05; ** = *p* < 0.01; *** = *p* < 0.001; **** = *p* < 0.0001).

### *C. jejuni*-mediated UPR activation is independent of NOD1 activity

To gain further insight into the relationship between *C. jejuni*-induced inflammation and UPR activation, we next explored if NOD1 activity is linked to *C. jejuni*-mediated UPR activation. Pre-treatment with ML130, a NOD1 inhibitor, did not affect *C. jejuni*- or thapsigargin-induced *CHOP* and spliced *XBP1* expression in either T84 or Caco-2 cells suggesting *C. jejuni*- and thapsigargin-induced UPR are independent of NOD1 activity (Fig 7A-D). Investigation of the impact of ML130 on *C. jejuni* and thpaisgargin-mediated IL-8 induction from T84 cells was further conducted. T84 cells treated with ML130 exhibited a significant reduction in thapsigargin-mediated IL-8 secretion compared to the untreated control suggesting thapsigargin-induced IL-8 is dependent on NOD1 activity (Fig 7E). In contrast, ML130 pre-treatment significantly increased *C. jejuni*-mediated IL-8 secretion from T84 cells (Fig 7E). Viability of *C. jejuni* and T84 cells was unaffected with ML130 pre-treatment (S3 Fig and S4 Fig). These results implicate that *C. jejuni* infection involving the combined effects of different virulence determinants results in an additional multifactorial response compared to thapsigargin treatment.

**Fig 7.**
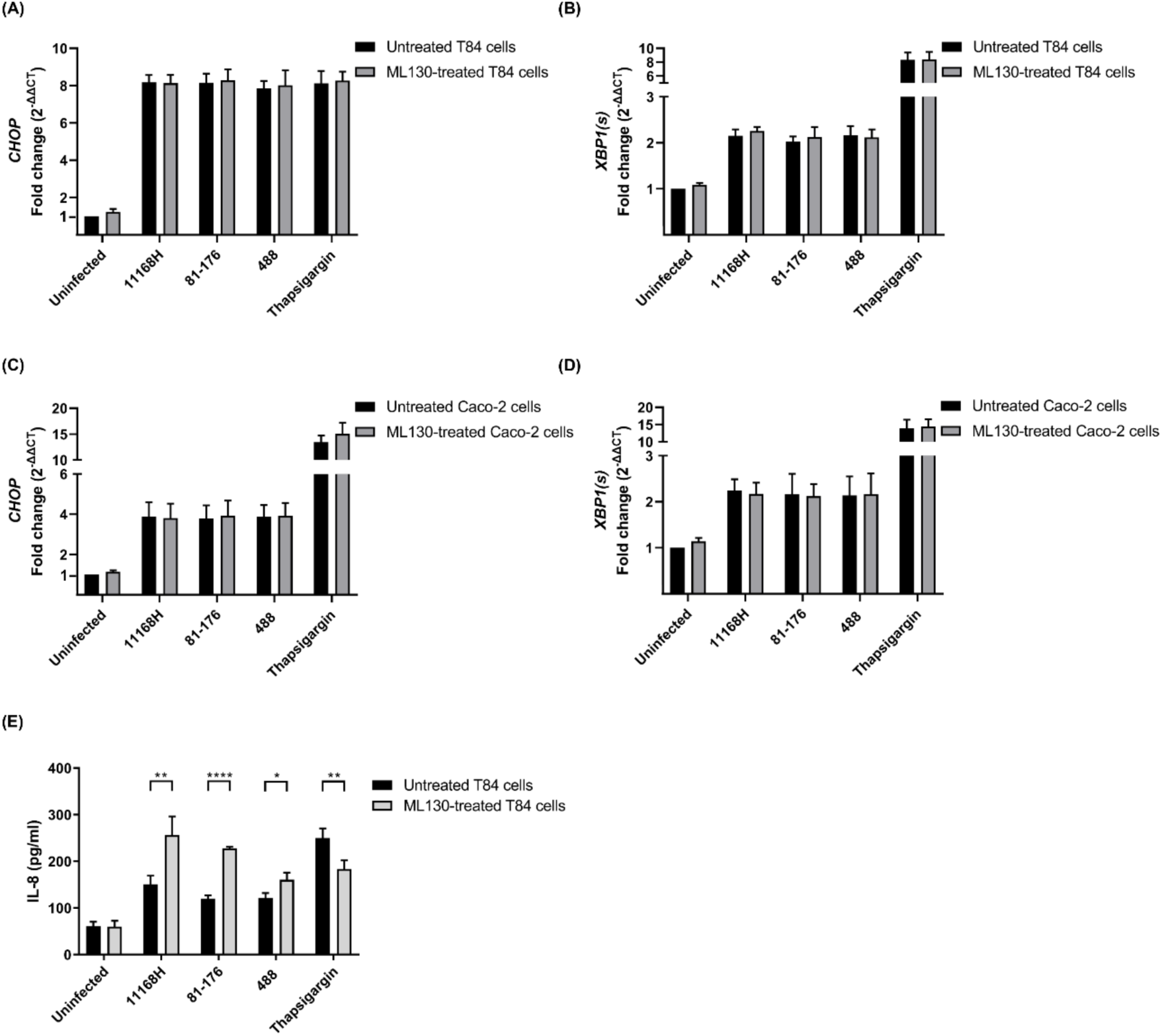
UPR-related gene expression in T84 and Caco-2 cells treated with the NOD1 inhibitor ML130 followed by *C. jejuni* infection or thapsigargin treatment and the impact of the NOD1 inhibitor ML130 on IL-8 release from T84 cells infected with *C. jejuni* or treated with thapsigargin. qRT-PCR showing expression of human (A, C) *CHOP* and (B, D) spliced *XBP1* [*XBP1(s)*] in T84 or Caco-2 cells infected with either *C. jejuni* 11168H, 81-176 or 488 wild-type strains (MOI of 200:1) or treated with 2 µM of thapsigargin for 24 h at 37°C in a 5% CO_2_ atmosphere. Prior to *C. jejuni* infection or thapsigargin treatment, T84 and Caco-2 cells were treated with 30 µM of ML130 for 4 h at 37°C in a 5% CO_2_ incubator. T84 and Caco-2 cells were further treated with 30 µM of ML130 during *C. jejuni* infection or thapsigargin treatment. *GAPDH* was used as an internal control. (E) T84 cells were pre-treated with 30 µM of ML130 for 4 h at 37°C in a 5% CO_2_ incubator. T84 cells were then infected with *C. jejuni* 11168H, 81-176 or 488 wild-type strains (MOI of 200:1) or treated with 2 µM thapsigargin for 24 h at 37°C in a 5% CO_2_ incubator. T84 cells were further treated with 30 µM of ML130 for 4 h during *C. jejuni* infection. Medium from each well was subjected to human IL-8 ELISA to measure the concentrations of IL-8. One sample *t*-test was performed. Asterisks denote a statistically significant difference (* = *p* < 0.05; ** = *p* < 0.01; **** = *p* < 0.0001).

### *C. jejuni*-mediated UPR activation is independent of Ca^2+^ release

Interruption of Ca^2+^ homeostasis by thapsigargin induces ER stress and activates the UPR. Thapsigargin inhibits SERCA which transports Ca^2+^ from the cytoplasm into the ER and consequently increases intracellular free Ca^2+^ within the cytosol [25]. The concentration of intracellular free Ca^2+^ within the cytosol in T84 cells infected with *C. jejuni* or thapsigargin was investigated (Fig 8A). All three *C. jejuni* wild-type strains and thapsigargin significantly increased the concentration of intracellular free Ca^2+^ within the cytosol compared to uninfected or untreated T84 cells (Fig 8A). In addition, T84 cells were treated with BAPTA-AM prior to *C. jejuni* infection or thapsigargin treatment. BAPTA-AM pre-treatment significantly reduced the *C. jejuni*- or thapsigargin-mediated increases in intracellular Ca^2+^ within the cytosol (Fig 8A). Viability of *C. jejuni* and T84 cells was unaffected with BAPTA-AM pre-treatment (S3 Fig and S4 Fig).

**Fig 8.**
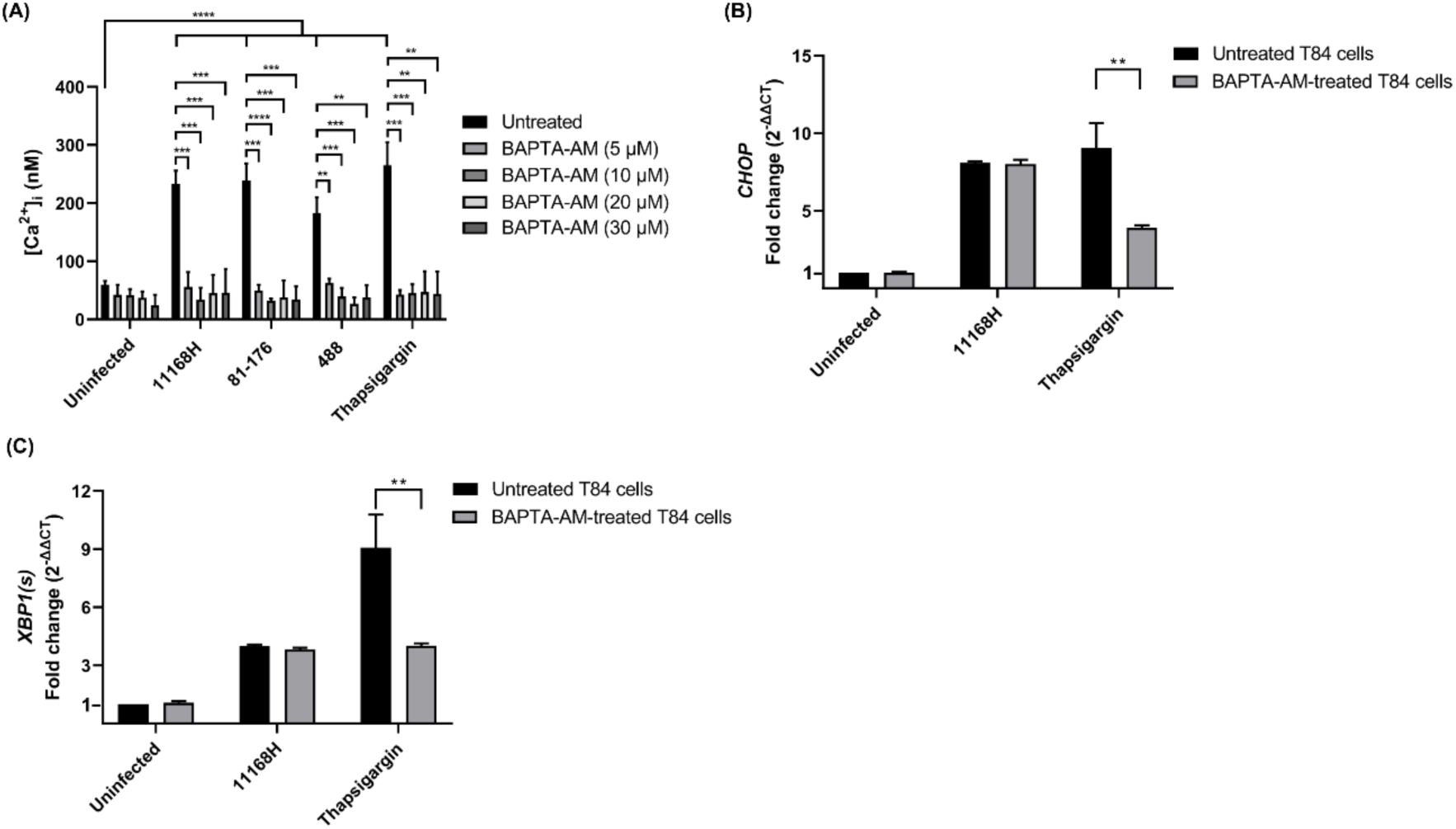
The effect of the Ca^2+^ chelator BAPTA-AM on the increases in intracellular Ca^2+^ resulting from *C. jejuni* infection or thapsigargin treatment in human IECs and UPR-related gene expression in T84 cells pre-treated with BAPTA-AM followed by *C. jejuni* infection or thapsigargin treatment. T84 cells grown in a 96-well plate were pre-treated with 5, 10, 20 or 30 µM of BAPTA-AM for 1 h at 37°C in a 5% CO_2_ incubator. T84 cells were then infected with *C. jejuni* 11168H, 81-176 or 488 wild-type strains (MOI of 200:1) or treated with 2 µM thapsigargin for 24 h at 37°C in a 5% CO_2_ incubator. After infection or treatment, 5 μM of cell-permeable dye Fura-2, AM was used to detect Ca^2+^. The two sets of fluorescence with 340 nm excitation and 510 nm emission (λ_1_) and 380 nm excitation and 510 nm emission (λ_2_) were recorded and intracellular Ca^2+^ concentration was calculated [26]. qRT-PCR showing expression of human (B) *CHOP* and (C) spliced *XBP1* [*XBP1(s)*] in T84 cells infected with either *C. jejuni* 11168H wild-type strain (MOI of 200:1) or treated with 2 µM of thapsigargin for 24 h at 37°C in a 5% CO_2_ atmosphere. Prior to *C. jejuni* infection or thapsigargin treatment, T84 cells were pre-treated with 10 µM of BAPTA-AM for 1 h at 37°C in a 5% CO_2_ incubator. *GAPDH* was used as an internal control. One sample *t*-test was performed. Asterisks denote a statistically significant difference (** = *p* < 0.01; *** = *p* < 0.001; **** = *p* < 0.0001).

Having established that *C. jejuni* increases intracellular free Ca^2+^ within the cytosol in human IECs, we hypothesised that *C. jejuni*-mediated UPR activation may be associated with the increase in intracellular Ca^2+^ within the cytosol. While pre-treatment with BAPTA-AM significantly lessened thapsigargin-mediated upregulation of *CHOP* and spliced *XBP1* in T84 cells, *C. jejuni*-mediated upregulation of *CHOP* and spliced *XBP1* was unaffected (Fig 8B and 8C). Collectively, these observations indicate that *C. jejuni*-mediated UPR activation is independent of any disruption of intracellular Ca^2+^ homeostasis.

### *C. jejuni* capsule and flagella mutants exhibit selective impairment in PERK pathway activation

We next explored *C. jejuni* virulence determinants which are responsible for activation of the PERK and IRE1α pathways in human IECs. Intriguingly, infection with the 11168H *kpsM* and *flaA* mutants resulted in significantly reduced upregulation of *CHOP* in T84 cells compared to the 11168H wild-type strain (Fig 9A). However, none of these mutants including the *kpsM* and *flaA* mutants resulted in significant differences in expression of spliced *XBP1* compared to the wild-type strain (Figs 9B and 9C). The heat-killed 11168H wild-type strain was significantly less upregulated for both *CHOP* and spliced *XBP1* expression compared to the live 11168H wild-type strain (Figs 9A and 9B). Growth kinetics of *C. jejuni* 11168H wild-type strain and different mutants (S5 Fig) and abilities of *C. jejuni* interaction with and invasion of T84 cells were investigated to query if *C. jejuni*-mediated UPR activation is related to bacterial interaction and invasion to human IECs (S6 Fig). The 11168H *cdtABC* operon mutant exhibited significantly enhanced interactions compared to the wild-type strain but did not exhibit a significant difference in invasion (S6 Fig). The 11168H *cdtA*, *htrA*, *kpsM* and *flaA* mutants showed significant reductions in both adhesion to and invasion of T84 cells compared to the wild-type strain. These observations suggest that *C. jejuni* capsule- and flagella-associated adhesion and invasion might be responsible for *C. jejuni*-mediated UPR activation in human IECs. Furthermore, these results indicate that different *C. jejuni* virulence determinants are differentially involved in activation of the branches of the UPR.

**Fig 9.**
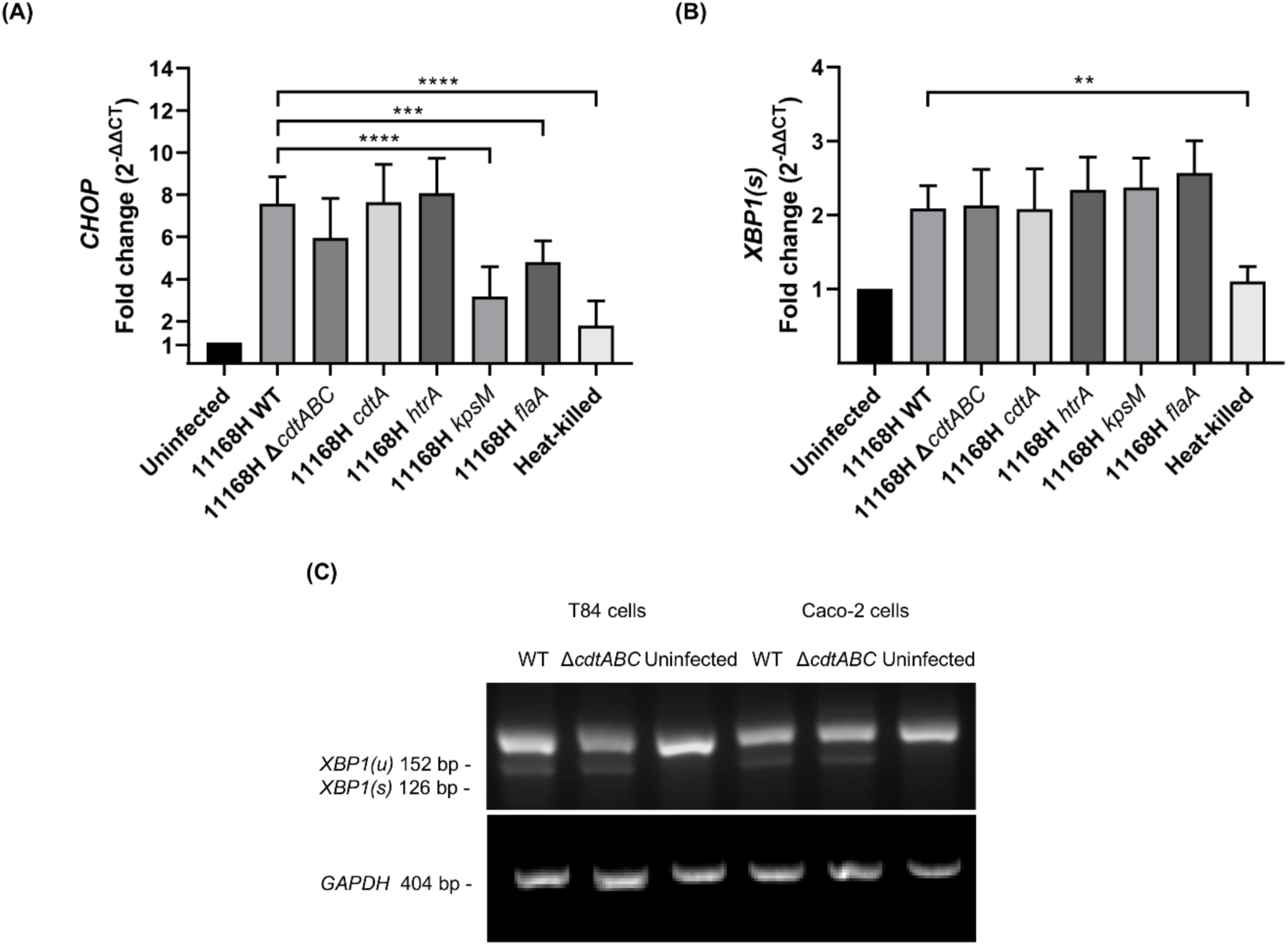
UPR-related gene expression following *C. jejuni* infection. qRT-PCR showing expression of human (A) *CHOP* and (B) spliced *XBP1* [*XBP1(s)*] in T84 cells infected with either *C. jejuni* 11168H wild-type strain, mutants or heat-killed 11168H wild-type strain for 24 h at 37°C in a 5% CO_2_ atmosphere (MOI of 200:1). (C) RT-PCR showing expression of unspliced and spliced *XBP1* [*XBP1(u)* and *XBP1(s)*] in T84 and Caco-2 cells infected with either *C. jejuni* 11168H wild-type strain or *cdtABC* operon mutant for 24 h at 37°C in a 5% CO_2_ atmosphere. *GAPDH* was used as an internal control. One sample *t*-test was performed. Asterisks denote a statistically significant difference (** = *p* < 0.01; *** = *p* < 0.001; **** = *p* < 0.0001).

## Discussion

The UPR is a pivotal component of host cell ER homeostasis involved in the response to ER stress [4]. As the UPR can be activated as a host defence mechanism against invading microbial pathogens, UPR activation is closely implicated with proinflammatory signalling [4]. However, the UPR can also be activated by some pathogenic microorganisms for survival purposes [4]. The link between *C. jejuni*-mediated inflammation and the UPR remains unclear. Compared to other enteric pathogens, *C. jejuni*-mediated UPR activation and its downstream consequences remains to be explored. This is the first study that implicates *C. jejuni*-mediated inflammation and its association with UPR activation in human IECs. Initially, we identified and examined the ability of three different *C. jejuni* strains to modulate each UPR pathway in human IECs *in vitro*. A previous study showed that *C. jejuni* induced only the PERK pathway in Caco-2 cells [24]. Intriguingly, in this study we demonstrated that *C. jejuni* 11168H, 81-176 and 488 strains induced the PERK and IRE1α pathways in both T84 and Caco-2 cells, whereas only 81-176 activated the ATF6 pathway in T84 cells but not in Caco-2 cells. These differing observations may be due to differences in experimental conditions.

Data presented in this study suggest that *C. jejuni* induces the ATF6 pathway in a cell line- and strain-dependent manner. T84 and Caco-2 cells possess distinct cellular characteristics which might be responsible for such differential ATF6 activation upon *C. jejuni* infection. T84 cells exhibit characteristics of enterocytes and express higher levels of TLR4, whereas Caco-2 cells exhibit characteristics of colonocytes and express higher levels of TLR5 [27]. Variation in ATF6 activation upon *C. jejuni* infection may be reflected by differential host-pathogen interactions in the two cell-lines which possess distinct characteristics.

In addition, we hypothesise that 81-176 might possess additional bacterial determinants (the pTet and pVir plasmids which encode putative Type IV secretion system (T4SS)) which may induce the ATF6 pathway in T84 cells [28, 29]. The activation of ATF6 observed only in IECs infected with 81-176 suggests that the activation of the UPR induced by this pathogen is not attributable to universal factors like the disruption of Ca^2+^ homeostasis. This finding indicates that the specific mechanisms underlying UPR activation in this context may be distinct from those typically associated with other stressors that broadly induce the UPR.

As UPR activation can affect microbial fitness in host cells, some bacterial pathogens can modulate the UPR upon infection [4]. These UPR-activating bacteria such as *Salmonella enterica*, *Brucella melitensis* and *Chlamydia* species can take advantage of UPR activation for intracellular replication [21–23]. In contrast, as the UPR plays a role in bacterial clearance via crosstalk between UPR signalling and innate immune response [4, 6, 30], certain microbial pathogens such as *Legionella pneumophila* dampen UPR activation to evade host surveillance system [20, 31]. The impact of UPR activation on *C. jejuni* adhesion, invasion and intracellular survival in human IECs remains unclear. Surprisingly, chemical pre-activation of all three UPR pathways using thapsigargin significantly impaired intracellular survival of *C. jejuni* within human IECs. Similarly, selective inhibition of either the PERK or IRE1α pathway by different UPR inhibitors resulted in significantly increased numbers of intracellular *C. jejuni* within human IECs. These results indicate each UPR pathway exhibits anti-bacterial effects on *C. jejuni* and the UPR is activated as a host defence mechanism. In addition, the observations suggest that *C. jejuni* does not take advantages of the UPR for its replication. Unlike other intracellular bacteria, which can utilise or suppress UPR activation for replication within host cells, *C. jejuni* does not appear to replicate within human IECs but can survive within the host cell for more than 24 hours [32]. Given that the UPR is activated upon *C. jejuni* infection and the UPR limits the number of intracellular *C. jejuni* within human IECs, it is logical to speculate that the *C. jejuni* may not be able to subvert the UPR and subsequent UPR-mediated inflammation may interrupt *C. jejuni* replication and survival for extended periods of time within human IECs. However, it is also possible that UPR activation and inflammation may potentially contribute to downstream *C. jejuni*-mediated damage.

As briefly mentioned above, the UPR is closely linked to proinflammation [6]. We also demonstrated that *C. jejuni*-induced inflammation in human IECs is correlated with UPR activation. Inhibition of the IRE1α and PERK pathways by KIRA6 and GSK2656157 significantly reduced *C. jejuni*- or thapsigargin-mediated IL-8 secretion. This observation suggests these two UPR pathways are involved in *C. jejuni*- and thapsigargin-mediated inflammation. In innate immune signalling, tumour necrosis factor receptor associated factor (TRAF2) interacts with phosphorylated IRE1α and induces inflammation by activating c-Jun N-terminal kinase (JNK) and NF-κB signalling pathways [33]. Given that STF-083010 inhibits only RNase activity but not kinase activity of IRE1α [34], the reason why no difference in thapsigargin-mediated IL-8 secretion in T84 cells treated with STF-083010 may be because IRE1α can still undergo autophosphorylation and can be recognised by TRAF2 resulting in inflammation. In other words, thapsigargin-mediated inflammation is dependent on the kinase activity of IRE1α rather than RNase activity which is supported by a previous study [35]. Surprisingly, we showed STF-083010 increased *C. jejuni*-mediated IL-8 secretion. There might be additional cellular effects of STF-083010 on immune signalling upon *C. jejuni* infection. In addition, it is possible that increased numbers of intracellular *C. jejuni* resulted from STF-083010 pre-treatment might lead to elevated *C. jejuni*-induced IL-8 secretion. Unlike KIRA6 and GSK2656157, STF-083010 may produce an environment where a correlation between bacterial burden within host cells and IL-8 induction is significant.

As non-canonical signalling of nucleotide-binding oligomerization domain 1 (NOD1) is involved in IRE1α-mediated inflammation [35, 36], we investigated if *C. jejuni*-mediated UPR activation and inflammation are associated with NOD1 activity. Intriguingly, we demonstrated that NOD1 activity is not required for *C. jejuni*- and thapsigargin-mediated UPR activation in human IECs. Our data is consistent with a previous study which showed there are no significant differences in thapsigargin-stimulated UPR activation in bone marrow-derived macrophages (BMDM) between NOD1/2^-/-^ and wild-type mice [35]. In contrast, ML130, a NOD1 inhibitor, showed differential effects on *C. jejuni*- and thapsigargin-mediated IL-8 secretion. NOD1 inhibition significantly reduced thapsigargin-mediated IL-8 secretion but significantly increased *C. jejuni*-mediated IL-8 secretion. This observation also supports the concept that thapsigargin-induced inflammation is dependent on kinase activity of IRE1α so that TRAF2 recognises phosphorylated IRE1α and further interacts with NOD1 for inflammatory response [35, 36].

In addition, our data suggests *C. jejuni*-mediated IL-8 secretion is correlated with UPR activation, but not associated with NOD1 activity. In contrast to our data which showed NOD1 inhibition resulted in increased *C. jejuni*-induced IL-8 expression, a previous study showed that NOD1 siRNA transfected cells exhibited reduced *C. jejuni*-induced IL-8 expression in Caco-2 cells compared to negative controls [19]. Such differing data in IL-8 secretion may be because of differences in the consequent physiological conditions which NOD1 siRNA and ML130 pre-treatment produce. ML130 directly interacts with NOD1 and may induce conformational changes which then interrupt intracellular trafficking of NOD1 [37]. ML130 pre-treatment also increases the localisation of NOD1 near the plasma membrane and reduces recruitment of a downstream adapter protein, receptor-interacting protein kinase (RIPK2) [37]. Alteration of the localisation of NOD1 and RIPK2 might affect fine-tuned cellular immune system resulting in increased *C. jejuni*-mediated IL-8 secretion. We speculate that ML130 may increase NOD1 near the plasma membrane resulting in enhanced detection of invading *C. jejuni* in human IECs. As a previous study demonstrated that binding of NOD1 ligand synergistically facilitates TLR-induced IL-8 secretion [38], with ML130 pre-treatment, the enhanced detection of *C. jejuni* by NOD1 might synergistically facilitate TLR-mediated inflammation.

Given that *C. jejuni* increases cytosolic free Ca^2+^ in human IECs via an unknown mechanism [26], we examined whether *C. jejuni*-mediated UPR activation is due to the increase in cytosolic free Ca^2+^. Consistent with the previous study [26], all three *C. jejuni* strains significantly increased intracellular free Ca^2+^ in the cytosol at the similar level as thapsigargin-mediated increase in intracellular free Ca^2+^. BAPTA-AM pre-treatment significantly impaired thapsigargin-mediated activation of PERK and IRE1α pathways indicating Ca^2+^ release is responsible for thapsigargin-induced UPR. In contrast, *C. jejuni*-mediated UPR activation was independent of Ca^2+^ release. Our data noted that 11168H and 488 strains activate the PERK and IRE1α pathways but not the ATF6 pathway whereas thapsigargin activates all three UPR pathways. Collectively, our data suggests *C. jejuni* may utilise different mechanisms for UPR activation which are unlike the mechanism of thapsigargin-mediated UPR. In addition, we hypothesise that *C. jejuni*-mediated UPR activation may not be due to general stimuli which induce activation of all three UPR pathways as Ca^2+^ reduction raises the amount of unfolded proteins within the ER lumen and generally activates all three UPR pathways [39]. It is also possible that *C. jejuni* may activate upstream cellular pathways directly or indirectly linked to each UPR pathway which is to be subject to further investigation.

We investigated the impact of specific virulence determinants in relation to UPR activation. Compared to the 11168H wild-type strain, the *cdtABC* operon and *cdtA* mutants did not exhibit any differences in activation of either the PERK or IRE1α pathways. Given that *C. jejuni* adhesion and invasion can facilitate IL-8 secretion which is independent of CDT activity and proinflammation is closely related to the UPR [6, 40, 41], a potential reason why no differences in UPR activation was observed between the *cdt* mutants and the wild-type strain may be because the UPR is still activated via *C. jejuni* adhesion and invasion. It may be possible that secretion of other virulence determinants such as outer membrane vesicles (OMVs) may be involved in UPR activation [14, 42]. A heterotrimeric AB-type CDT appears to require the ER-associated degradation (ERAD) pathway for retrotranslocation and intoxication and the UPR upregulates components of ERAD pathway [5, 43]. The observation that *C. jejuni* CDT did not activate the UPR indicate that retrotranslocation of *C. jejuni* CDT may utilise basal level of ERAD components without upregulation of ERAD components via UPR activation.

This study also demonstrated that *C. jejuni* capsular polysaccharide and flagella are associated with activation of the PERK pathway but not IRE1α pathways. This observation reemphasises the possibility that *C. jejuni* adherence and invasion might be responsible for activation of the PERK pathway. The results also indicate that effector proteins delivered through *C. jejuni* flagella may play a role in activating the UPR. *C. jejuni* CiaD, which is one of the Cia (*Campylobacter* invasion adhesion) proteins, has been shown to be involved in bacterial invasion into host cells via interaction with a host cell component called IQGAP1 (a Ras GTPase-activating-like protein) [44]. Further investigation into the effect of CiaD on UPR activation will clarify the mechanisms by which *C. jejuni* adherence and invasion induce the UPR. Intriguingly, none of the mutants used in this study showed any significant differences in IRE1α pathway activation implying different, as yet undetermined *C. jejuni* virulence determinants are involved in the activation of each UPR pathway. A previous study demonstrated that TLR2 and TLR4 activate the IREα pathway in the absence of ER stress inducers [45]. *C. jejuni* lipoproteins which are TLR2 and TLR4 agonists might be responsible for *C. jejuni*-mediated IREα pathway activation.

Our results reveal host cell UPR plays an important role in inflammation upon *C. jejuni* infection. While UPR activation can help eliminate the pathogen, its role in inducing inflammation might create a double-edged sword. This inflammation can lead to tissue damage, complicating the host’s response and aiding bacterial translocation across IECs. Therefore, it highlights the balance that the immune system must maintain between effectively combating infections and minimising damage to host tissues. Further investigation into how UPR-mediated inflammation can be modulated could provide insights into potential therapeutic approaches. Inhibiting UPR activation could serve as a potential therapeutic strategy for irritable bowel syndrome caused by *C. jejuni*, as well as other inflammatory conditions. We also provide new data implicating virulence determinants with key roles in contributing to UPR activation and that *C. jejuni*-induced UPR activation is independent of Ca^2+^ release within the cytosol. Further characterisation of crosstalk between *C. jejuni*-mediated UPR activation and inflammation and of potential molecular mechanisms of *C. jejuni*-mediated UPR activation will provide new insights into cellular mechanisms of *C. jejuni*-induced diarrhoeal disease.

## Materials and Methods

### Bacterial strains and growth conditions

*C. jejuni* wild-type strains used in this study are listed in S1 Table. For general growth, all *C. jejuni* strains were grown on Columbia Blood Agar (CBA) plates (Oxoid, UK) supplemented with 7% (v/v) horse blood in Alsever’s (TCS Microbiology, UK) and *Campylobacter* selective supplement Skirrow (Oxoid) or in *Brucella* broth (BD Diagnostics, UK) at 37 °C under microaerobic conditions (10% CO_2_, 5% O_2_ and 85% N_2_) (Don Whitley Scientific, UK). *E. coli* DH5α™ Competent Cells (New England BioLabs, UK) were grown on lysogeny broth (LB; Oxoid) agar plates or in LB broth at 37°C under aerobic conditions shaking at 200 rpm (Sanyo, UK). For growth of *C. jejuni* mutants, kanamycin (50 μg/ml) or chloramphenicol (10 μg/ml) were supplemented to CBA. If necessary, *E. coli* cultures were added with ampicillin (100 μg/ml) and chloramphenicol (50 μg/ml).

### Human intestinal epithelial cell culture

Human carcinoma cell line T84 cells (ECACC 88021101) and Caco-2 cells (ECACC 86010202) were obtained from European Collection of Authenticated Cell Cultures (ECACC). T84 and Caco-2 cells were cultured in complete growth medium which is composed of 1:1 mixture of Dulbecco’s modified Eagle’s medium and Ham’s F-12 medium (DMEM/F-12; Thermo Fisher Scientific, USA) with 10% Fetal Bovine Serum (FBS; Labtech, U.K), 1% non-essential amino acid (Sigma-Aldrich, USA) and 1% penicillin-streptomycin (Sigma-Aldrich). For the co-culture of human IECs with *C. jejuni*, complete growth medium without penicillin-streptomycin was used. Complete growth medium without penicillin-streptomycin and phenol red was used for the intracellular Ca^2+^ measurement assay. Cell lines were cultured at 37°C in a 5% CO_2_ humidified incubator.

### T84 and Caco-2 cells infection assays

Approximately 10^4^, 10^5^ or 2 x 10^5^ human IECs were seeded into each well of 96-, 24- or 6-well plates respectively. 96-well plates were then incubated overnight at 37°C in a 5% CO_2_ incubator to allow cell attachment. 24- and 6-well plates were then incubated at 37°C in a 5% CO_2_ incubator for 7 days prior to co-incubation with *C. jejuni.* Prior to the infection, human IECs were washed with pre-warmed phosphate-buffered saline (PBS; Thermo Fisher Scientific) three times and complete growth medium without penicillin-streptomycin was added into each well. *C. jejuni* strains grown on 24-h CBA plates were resuspended in PBS and bacterial suspension with appropriate optical density at 600 nm (OD_600_) was then coincubated with IECs for indicated time periods giving a multiplicity of infection (MOI) of 200:1. In some experiments, human IECs were pre-treated with chemicals and/or further co-incubated with either *C. jejuni* or thapsigargin as depicted in S2 Table. After pre-treatment, IECs were washed three times with pre-warmed PBS.

### Real time-quantitative polymerase chain reaction (qRT-PCR) analysis

Eukaryotic RNA was extracted from infected and uninfected IECs using PureLink^TM^ RNA Mini Kit (Thermo Fisher Scientific) and contaminating DNA was removed from each RNA sample using TURBO DNA-free kit (Ambion, USA) according to manufacturer’s instructions. The concentration and purity of RNA samples were determined using a NanoDrop ND-1000 spectrophotometer (Thermo Fisher Scientific). For complementary DNA (cDNA) synthesis, 400 ng of each RNA sample was first mixed with 1 μl of random hexamers (50 ng/µl; Invitrogen, USA) and 1 μl of dNTP mix (10 mM; Invitrogen) and incubated at 65°C for 5 minutes and snap cooled on ice. The sample was then further incubated with 1 μl of SuperScript III RT (200 U/μl; Invitrogen), 1 μl of RNaseOUT (40 U/μl; Invitrogen), 2 μl of 0.1 M DTT (Invitrogen), 2 μl of 25 mM MgCl2 (Thermo Fisher Scientific) and 4 μl of 5X RT buffer (Invitrogen) at 25°C for 10 minutes followed by a 50 minute incubation at 50°C and a 5 minute incubation at 85°C. The sample was cooled on ice and incubated with RNase H (5U/μl; Thermo Fisher Scientific) at 37°C for 20 minutes. For qRT-PCR, each reaction had 10 µl of SYBR Green PCR Master Mix (Applied Biosystems, USA), 1 µl of primer (20 pmol), 1 µl of cDNA and 10 µl of DEPC-treated water (Invitrogen). The sequence of each primer is listed in S3 Table. All reactions were run in a minimum of three biological replicates, each in technical triplicate on an ABI-PRISM 7500 instrument (Applied Biosystems). *GAPDH* expression in the same sample was also analysed to normalize expression levels of all target genes. Relative expression changes were determined using the comparative threshold cycle (C_T_) method [46].

### Semi-quantitative reverse transcription (RT-PCR) analysis

Each PCR reaction had 25 μl of MyTaq Red Mix (Bioline, UK), 1 μl of forward primer (10 pmol), 1 μl of reverse primer (10 pmol) described in S3 Table, 1.5 μl of cDNA and 21.5 μl of DEPC-treated water. For PCR reactions Tetrad-2 Peltier thermal cycler (Bio-Rad, U.K) was used. One cycle of PCR programme performed as 95°C for 15 seconds after 2 minutes in the first cycle, annealing at 50°C for 20 seconds, and extension at 72°C for 30 seconds. Total 36 cycles were repeated. The PCR products were loaded on the 1% (w/v) agarose gel and the gel was running for 1 h at 120 V. For detection of XBP1 splicing, the PCR products were loaded on a 2% (w/v) agarose gel. The gel was visualised using an Azure c600 (Azure Biosystems, USA).

### Construction of *C. jejuni cdtABC* operon mutant

*C. jejuni* 11168H *cdtABC* operon mutant was constructed by using splicing by overlap extension PCR (SOE PCR) [47]. Briefly, the appropriate SOE PCR primers were designed to amplify the left fragment and right fragment of *C jejuni* 11168H *cdtABC* operon (S3 Table). The inner primers are partially complementary to each other, and the overlapping region contains a BamHI restriction site. The left and right fragments were then combined and amplified using the outside primers. The purified PCR products were ligated into a pGEM^®^-T Easy Vector (Promega, USA) and transformed into *E. coli* DH5α™ Competent Cells (New England BioLabs, UK). The resulting plasmid was digested with BamHI and chloramphenicol resistance cassette was inserted into the digested plasmid. *E. coli* was then transformed with the plasmid with chloramphenicol cassette. After confirmation of insertion and orientation of antibiotic cassette, the resulting plasmid was transformed into *C. jejuni* by electroporation. Successful construction of *C. jejuni cdtABC* operon mutant was confirmed by PCR, Sanger sequencing and qRT-PCR.

### Sodium dodecyl-sulfate polyacrylamide gel electrophoresis (SDS-PAGE) and Western blot analysis

After infection, IECs were washed three times with ice-cold PBS and lysed with cold RIPA lysis and extraction buffer (Thermo Fisher Scientific) with Halt™ Protease & Phosphatase Inhibitor Single-Use Cocktail (Thermo Fisher Scientific) and centrifuged for 20 minutes at 17,000 × g at 4°C. Protein concentration of each sample was measured using Pierce™ Bicinchoninic acid (BCA) Protein Assay Kit (Thermo Fisher Scientific) according to the manufacturer’s instructions. Samples were then diluted to a desired concentration in DEPC-treated water and 4X Laemmli sample buffer (Sigma-Aldrich) and incubated for 5 minutes at 95°C. 30 μg of protein samples were separated using 4-12% (w/v) NuPAGE™ Bis-Tris gel in 1X NuPAGE™ MES buffer or MOPS buffer (Invitrogen). Proteins were transferred from the gel to a nitrocellulose membrane using iBlot® 2 transfer stacks (Life Technologies, USA) and iBlot® Gel Transfer Device (Invitrogen). The membrane was then blocked with 1X PBS containing 2% (w/v) skimmed milk (Sigma-Aldrich). After blocking, the membrane was probed with primary antibodies overnight at 4°C as described previously [48]. The following primary antibodies were used; GAPDH (ab181602; Abcam); human CHOP (NB600-1335; Novis Biologicals); human XBP1 (PA5-27650; Thermo Fisher Scientific). After the incubation, blots were probed with either goat anti-rabbit IgG 800 or goat anti-mouse IgG 800 (LI-COR Biosciences, USA) and visualized on a LI-COR Odyssey Classic (LI-COR Biosciences).

### Adhesion, invasion and intracellular survival assay

Adherence, invasion and intracellular assays were performed as described previously [48]. T84 and Caco-2 cells seeded in a 24-well plate were washed three times with pre-warmed PBS and pre-treated with various chemicals as described above. Then IECs were infected with *C. jejuni* with OD_600_ 0.2 (MOI of 200:1) and incubated for 3 h at 37°C in a 5% CO_2_ incubator. For interaction (adhesion and invasion) assay, IECs were washed three times with pre-warmed PBS to remove unbound extracellular *C. jejuni* and then lysed with 0.1% (v/v) Triton X-100 (Sigma-Aldrich) for 20 minutes at room temperature. The cell lysates were serial-diluted and plated on to CBA plates. The CBA plates were incubated for 48 h at 37°C under microaerobic condition and the number of interacting bacteria was determined.

Invasion assay was performed by additional step of treatment of gentamicin (150 µg/ml) for 2 h to kill extracellular bacteria. After treatment, IECs were washed three times with PBS, lysed and plated on to CBA plates as described above. For intracellular survival assay, after infection with *C. jejuni*, IECs were treated with gentamicin (150 µg/ml) for 2 h to kill extracellular bacteria and further incubated with gentamicin (10 µg/ml) for 18 h. Cell lysis and inoculation on to CBA plates were performed as described above.

### Intracellular Ca^2+^ measurement assay

To measure intracellular Ca^2+^ concentration, T84 cells were seeded into a clear bottom and black 96-well plate and incubated overnight at 37°C in a 5% CO_2_ atmosphere. T84 cells were pre-treated with BAPTA-AM for 1 h prior to infection with *C. jejuni* or treatment with thapsigargin as described above. After treatment, T84 cells were infected with *C. jejuni* (MOI of 200:1) or treated with 2 μM of thapsigargin for 3 or 24 h at 37°C in a 5% CO_2_ atmosphere. IECs were then washed three times with 150 μl of HEPES buffered saline solution (BSS) (PromoCell). T84 cells were probed with cell-permeable dye Fura-2, AM (Invitrogen) to detect intracellular Ca^2+^. The dye solution contained 5 μM Fura-2, AM, 0.05% (v/v) Pluronic F-127 (Invitrogen) and 2.5 mM probenecid (Invitrogen), and HEPES BSS. T84 cells were incubated with 100 μl of dye solution for 1 h protected from light at 37°C in a 5% CO_2_ incubator. After 1 h incubation, T84 cells were washed twice with 200 μl of DMEM/F-12 without phenol red and incubated for additional 30 minutes at 37°C in a 5% CO_2_ incubator. At the end of the experiment, for the maximal fluorescence, 0.1% (v/v) Triton X-100 was added to lyse the cells and release intracellular Fura-2, AM from the cells. For the minimal fluorescence, 0.1% (v/v) Triton X-100 was added with subsequent addition of 30 μM BAPTA-AM. Then the two sets of fluorescence with 340 nm excitation and 510 nm emission (λ1) and 380 nm excitation and 510 nm emission (λ2) were recorded using a SpectraMax M3 Multi-Mode Microplate Reader. Intracellular Ca^2+^ concentration was calculated using the equation below [26].

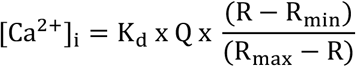

In the equation, R indicates the ratio of fluorescence intensity at λ_1_ and λ_2_. λ_1_ detects Ca^2+^-bound Fura-2 and λ_2_ detects Ca^2+^-free Fura-2. Q is the ratio of the minimal fluorescence intensity and the maximal fluorescence intensity at λ_2_. K_d_ is the dissociation of constant of the dye. The K_d_ value of Fura-2 at 37°C is 224 nM [49]. R_min_ is the ratio of minimal fluorescence intensity at λ_1_ and λ_2_ and R_max_ is the ratio of maximal fluorescence intensity at λ_1_ and λ_2_.

### *C. jejuni* viability test under experimental conditions

*C. jejuni* suspensions with OD_600_ 0.2 were treated with chemicals as described in S2 Table. After treatment, serial dilutions were performed and 10 µl of each dilution was spotted in triplicate on to CBA plates. The plates were incubated under microaerobic atmosphere at 37°C for 48 h. The CFUs from each spot were enumerated. Three biological replicates were performed for all experiments.

### Lactate dehydrogenase (LDH) cytotoxicity assay

According to the manufacturer’s instructions, CyQUANT™ LDH Cytotoxicity Assay kit (Thermo Scientific) was used to determine cytotoxicity of chemical treatments on IECs. Briefly, 10 µl of DEPC-treated water was added to one set of triplicate wells of IECs for spontaneous LDH activity controls. For chemical treated samples, chemicals were diluted in 10 µl of DEPC-treated water to give the final concentration as described in S2 Table. For maximum LDH activity controls, nothing was added at this step. The plate was incubated at 37°C in a 5% CO_2_ incubator for the indicated time as described in S2 Table. After treatment, 10 µl of 10X Lysis Buffer was added to the set of triplicate wells of maximum LDH activity controls and incubated at 37°C with 5% CO_2_ for 45 minutes. After incubation, 50 µl of medium from each well was transferred to a new clear bottom and black 96-well plate. Then 50 µl of reaction mixture was added to each well. The plate was then incubated at room temperature for 30 minutes protected from light. 50 µl of Stop Solution was then added to each well. The absorbance at both 490 nm and 680 nm was measured using SpectraMax M3 Multi-Mode Microplate Reader (Molecular Devices, USA). The LDH activity was determined by subtracting the absorbance at 680 nm from the absorbance at 490 nm. Cytotoxicity (%) was calculated as described in the manufacturer’s instructions.

### Statistical analysis and graphing

A minimum of three biological replicates were performed in all experiments. Three technical replicates were performed within each biological replicate. For statistical analysis and graphing, GraphPad Prism 8 for Windows (GraphPad Software, USA) was used. One sample *t*-test was performed to compare the mean of each data set with a hypothetical value 1. Unpaired *t*-test was performed to compare the mean between two independent data sets for significance with * indicating *p* < 0.05, ** indicating *p* < 0.01, *** indicating *p* < 0.001, and **** indicating *p* < 0.0001.

## Supporting information

**S1 Table.**
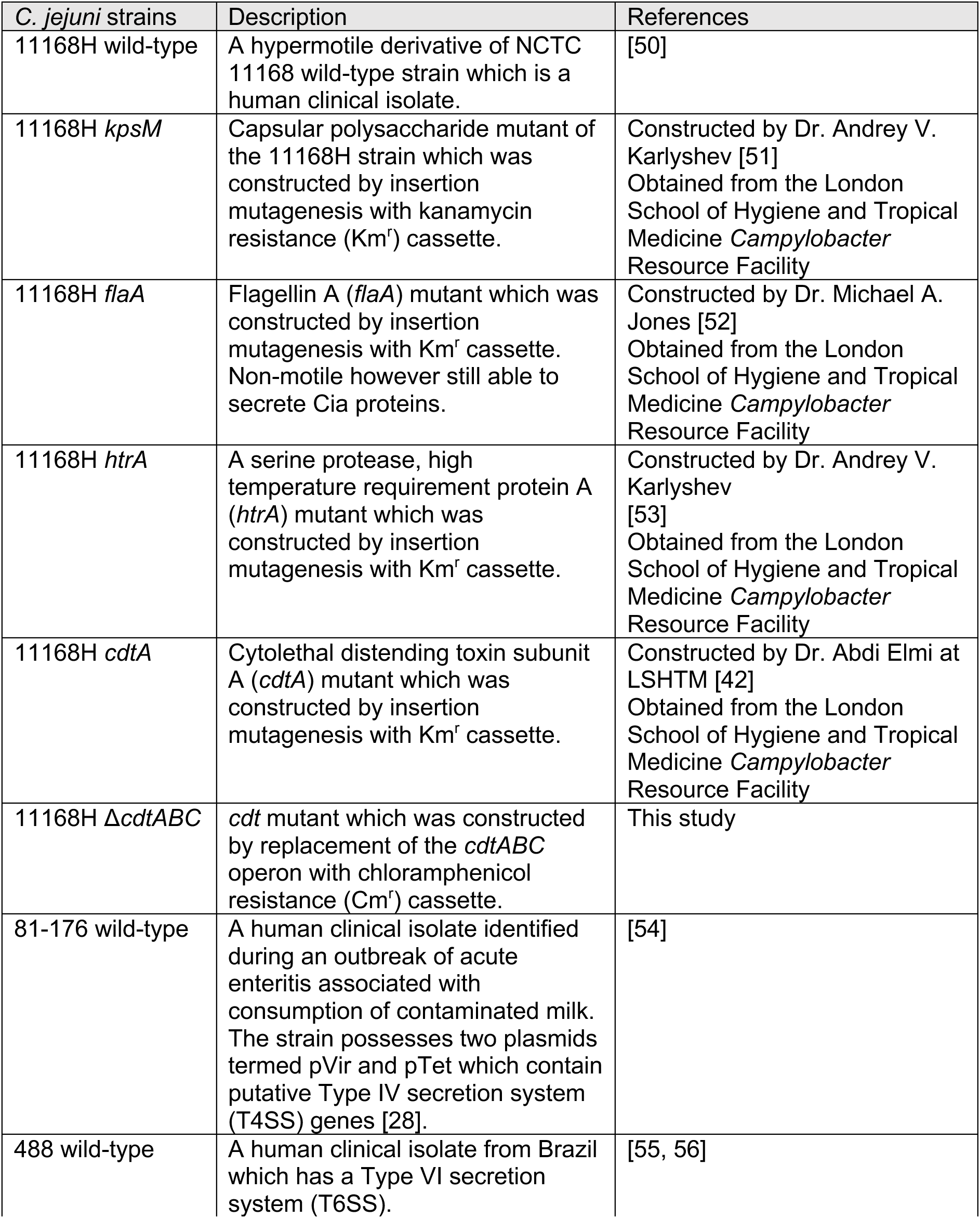
*C. jejuni* strains used in this study.

**S2 Table.** Chemicals and experimental conditions used in this study.

| Chemicals | Company | Description | Experimental condition |
| --- | --- | --- | --- |
| Thapsigargin (Tg) | Sigma-Aldrich | An inhibitor of sarco/endoplasmic reticulum Ca<sup>2+</sup>-ATPases (SERCA) | 2 μM for 6 h prior to infection |
|  |  | resulting in ER stress [25] |  |
| KIRA6 | Selleckchem (Germany) | IRE1α kinase and RNase inhibitor [34, 57] | 3 μM for 4 h pre-treatment before and during infection or Tg treatment |
| STF-083010 | Sigma-Aldrich | IRE1α RNase inhibitor [34] | 100 μM for 4 h pre-treatment before infection or Tg treatment |
| GSK2656157 | Selleckchem | ATP-competitive PERK inhibitor [58] | 3 μM for 4 h pre-treatment before infection or Tg treatment and during infection or Tg treatment |
| ML130 (Nodinitib-1) | Selleckchem | NOD1 inhibitor [37] | 30 μM for 1 h pre-treatment before infection or Tg treatment and during infection or Tg treatment |
| BAPTA-AM | Invitrogen | Cell-permeable Ca<sup>2+</sup> chelator [59] | 5, 10, 20, or 30 μM 1 h pre-treatment before infection or Tg treatment |

**S3 Table.**
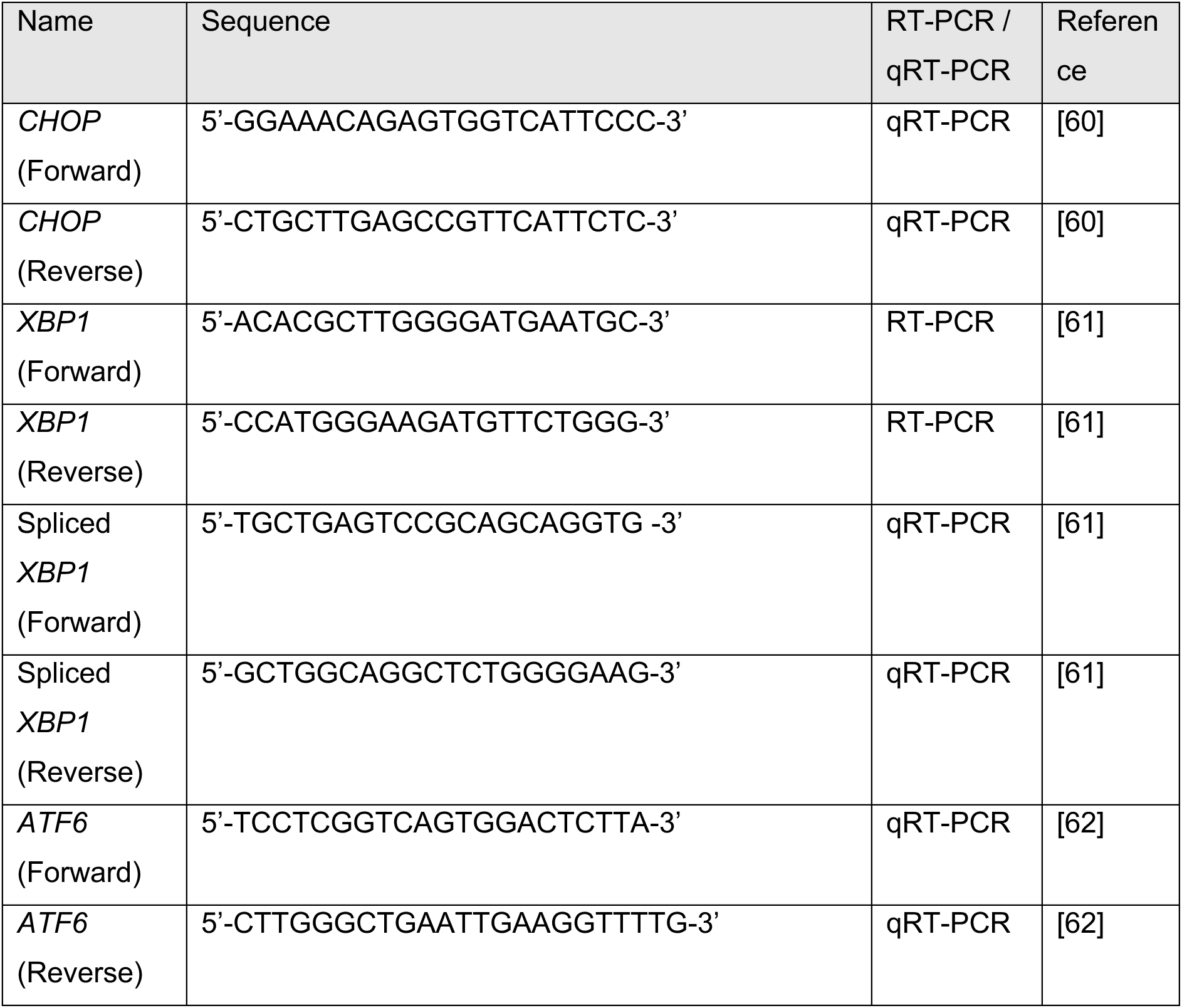
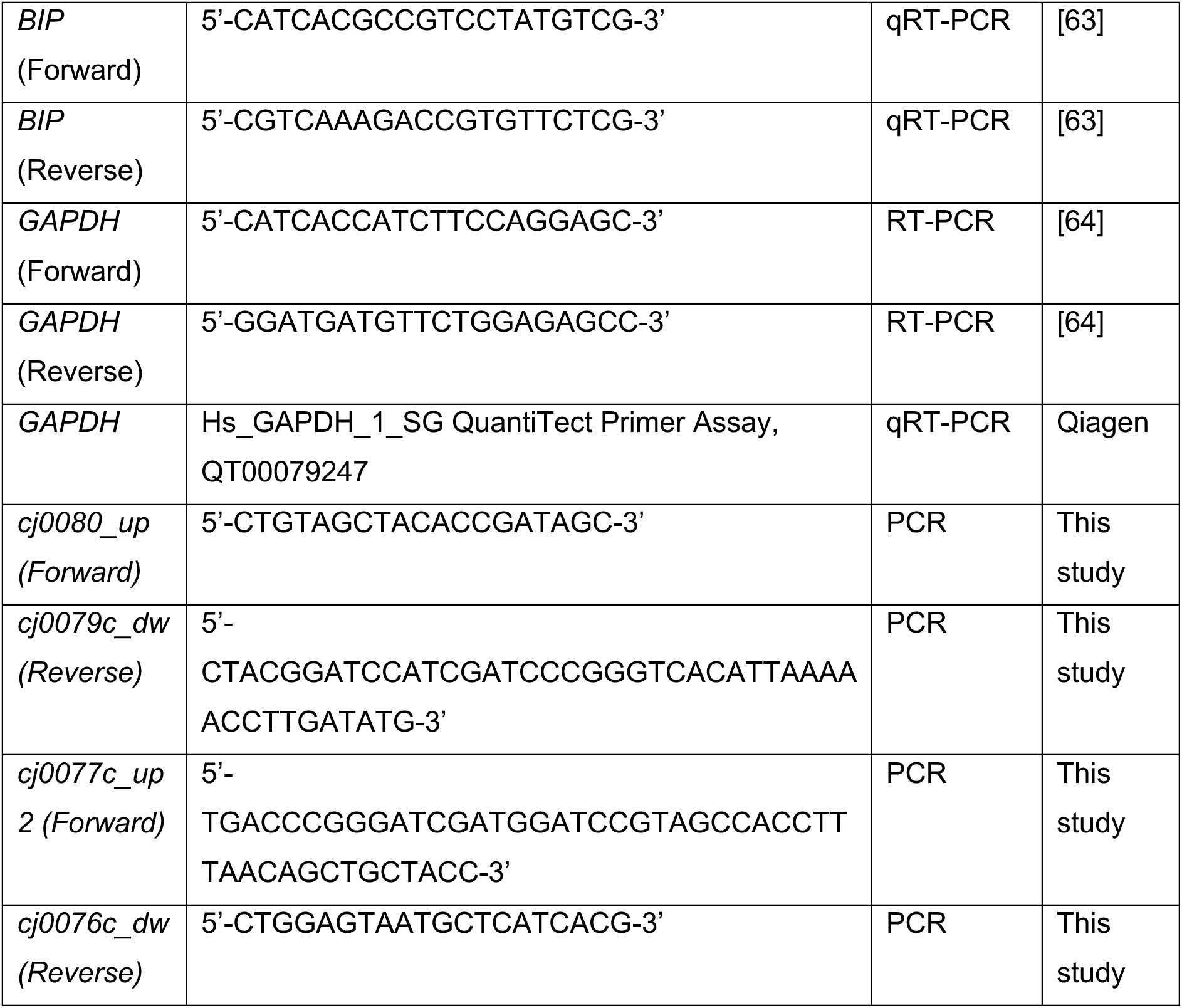
Primers used for RT-PCR and qRT-PCR.

**S1 Fig.**
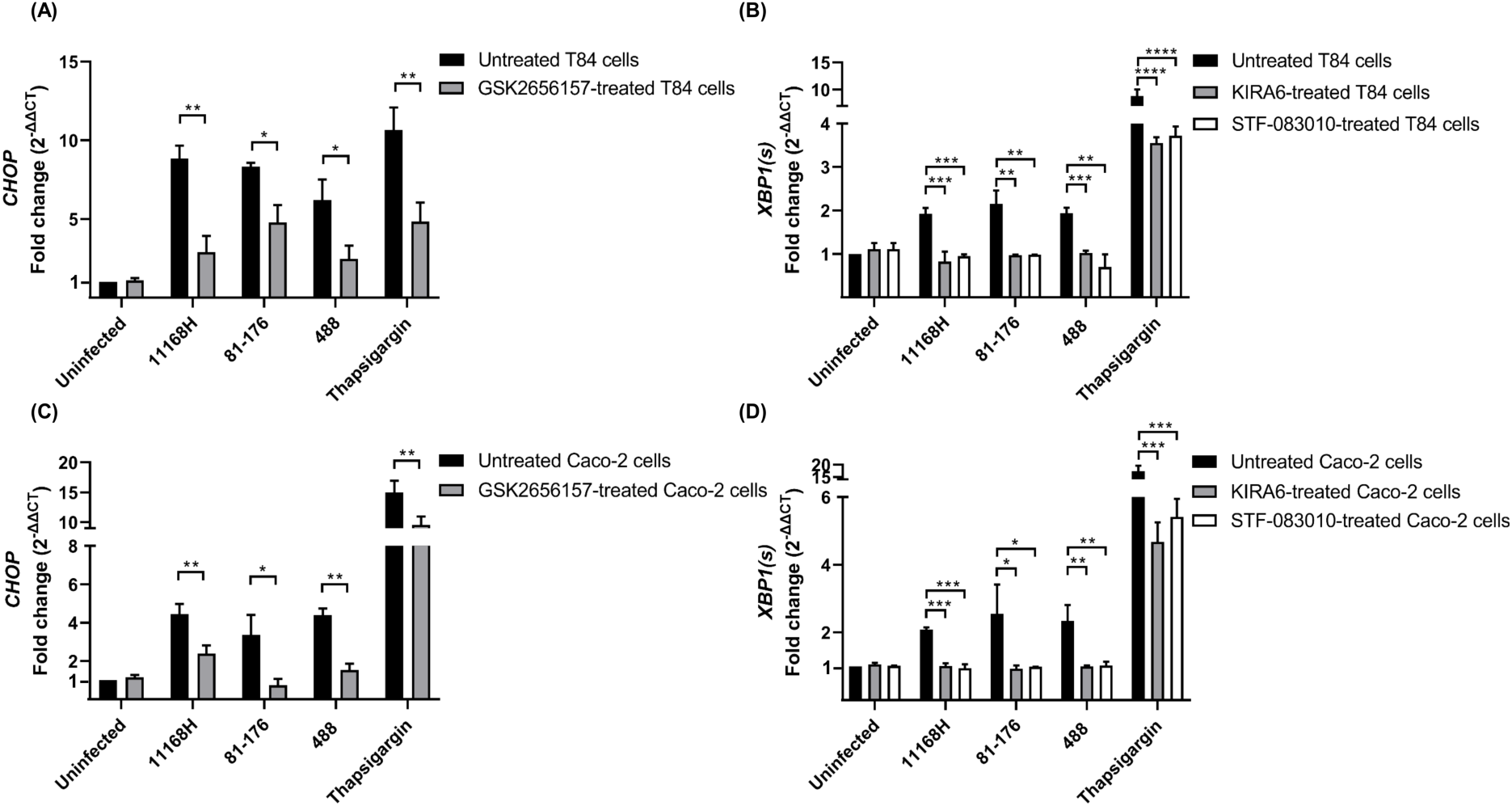
UPR-related gene expression in T84 and Caco-2 cells treated with UPR inhibitors followed by *C. jejuni* infection. qRT-PCR showing expression of human (A, C) *CHOP* and (B, D) spliced *XBP1* [*XBP1(s)*] in T84 or Caco-2 cells infected with either *C. jejuni* 11168H, 81-176 or 488 wild-type strains (MOI of 200:1) or treated with 2 µM of thapsigargin for 24 h at 37°C in a 5% CO_2_ atmosphere. Prior to *C. jejuni* infection or thapsigargin treatment, T84 and Caco-2 cells were treated with 3 µM of KIRA6, 100 µM of STF-083010 or 3 µM of GSK2656157 for 4 h at 37°C in a 5% CO_2_ incubator. T84 and Caco-2 cells were further treated with 3 µM of KIRA6 and 3 µM of GSK2656157 during *C. jejuni* infection or thapsigargin treatment. *GAPDH* was used as an internal control. Three biological and three technical replicates were performed for each experiment. Asterisks denote a statistically significant difference (* = *p* < 0.05; ** = *p* < 0.01; *** = *p* < 0.001; **** = *p* < 0.0001).

**S2 Fig.**
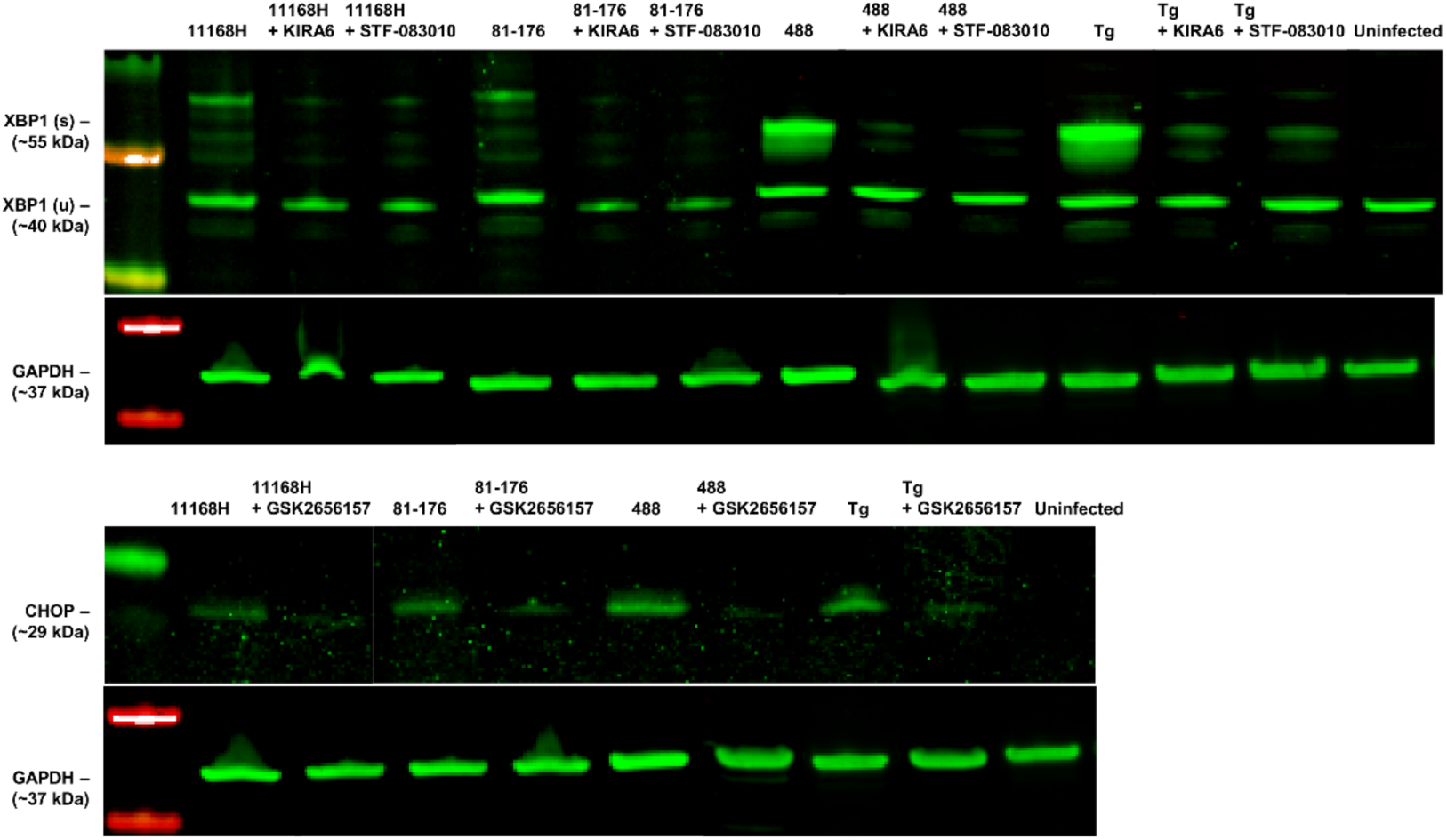
UPR-related protein expression in T84 cells treated with UPR inhibitors followed by *C. jejuni* infection. Western blotting showing protein levels of both spliced & unspliced forms of XBP1 [XBP1(u) & XBP1(s)] and CHOP in T84 cells infected with *C. jejuni* 11168H, 81-176 or 488 wild-type strains (MOI of 200:1) or treated with 2 µM of thapsigargin (Tg) for 24 h at 37°C in a 5% CO_2_ incubator. Prior to *C. jejuni* infection or thapsigargin treatment, T84 and Caco-2 cells were pre-treated with 3 µM of KIRA6, 100 µM of STF-083010 or 3 µM of GSK2656157 for 4 h at 37°C in a 5% CO_2_ incubator. T84 and Caco-2 cells were further treated with 3 µM of KIRA6 and 3 µM of GSK2656157 during *C. jejuni* infection or thapsigargin treatment. GAPDH was used as an internal control.

**S3 Fig.**
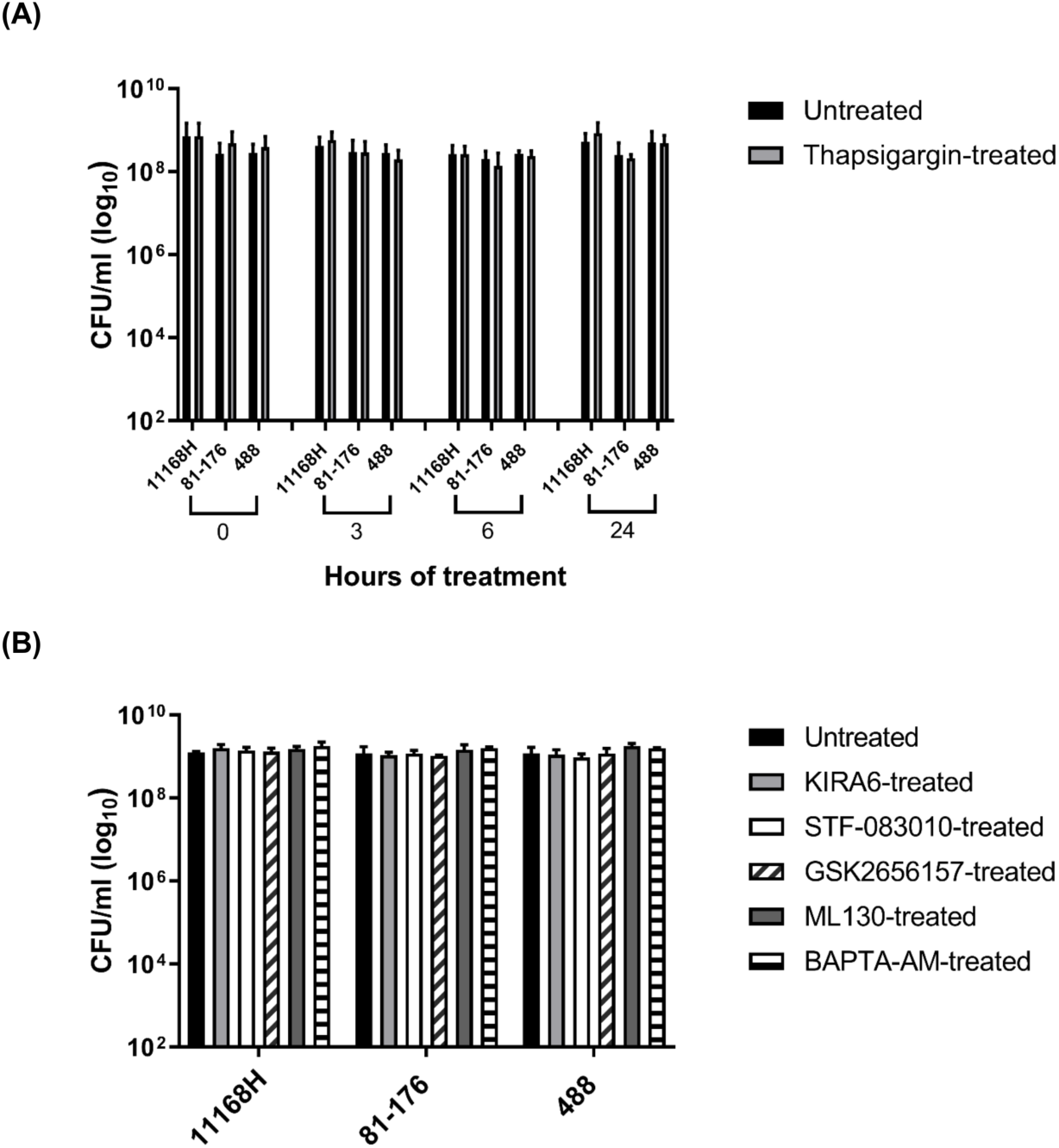
Assessment of *C. jejuni* vability with thapsigargin treatment. *C. jejuni* 11168H, 81-176 and 488 wild-type strains were treated with (A) 2 µM of thapsigargin for 3-, 6- or 24-h or (B) 3 µM of KIRA6, 3 µM of GSK2656157 or 30 µM of ML130 for 24 h or treated with 100 µM of STF-083010 for 4 h or 10 µM of BAPTA-AM for 1 h at 37°C in a 5% CO_2_ incubator. CFU/ml were recorded after incubation. Three biological and three technical replicates were performed for each experiment.

**S4 Fig.**
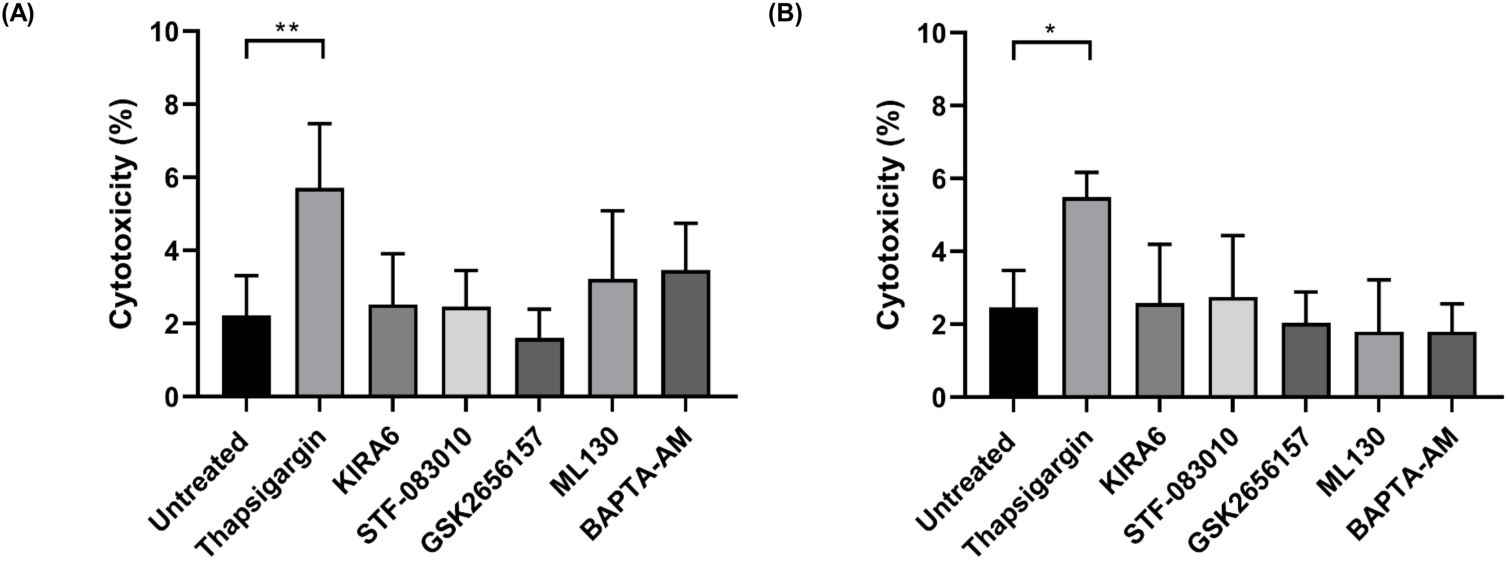
Measurement of cytotoxicity of chemical treatment on T84 and Caco-2 cells. (A) T84 or (B) Caco-2 cells grown in a 96-well plate were treated with 3 µM of KIRA6, 3 µM of GSK2656157 or 30 µM of ML130 for 24 h or treated with 2 µM of thapsigargin for 6 h, 100 µM of STF-083010 for 4 h or 10 µM of BAPTA-AM for 1 h at 37°C in a 5% CO_2_ incubator. Medium from each well was then analysed using a LDH assay. Cytotoxicity (%) was calculated by using the following equation: (*C. jejuni* infected LDH activity – spontaneous LDH activity) / (maximum LDH activity – spontaneous LDH activity) x 100. Three biological and three technical replicates were performed for each experiment. Asterisks denote a statistically significant difference (* = *p* < 0.05; ** = *p* < 0.01).

**S5 Fig.**
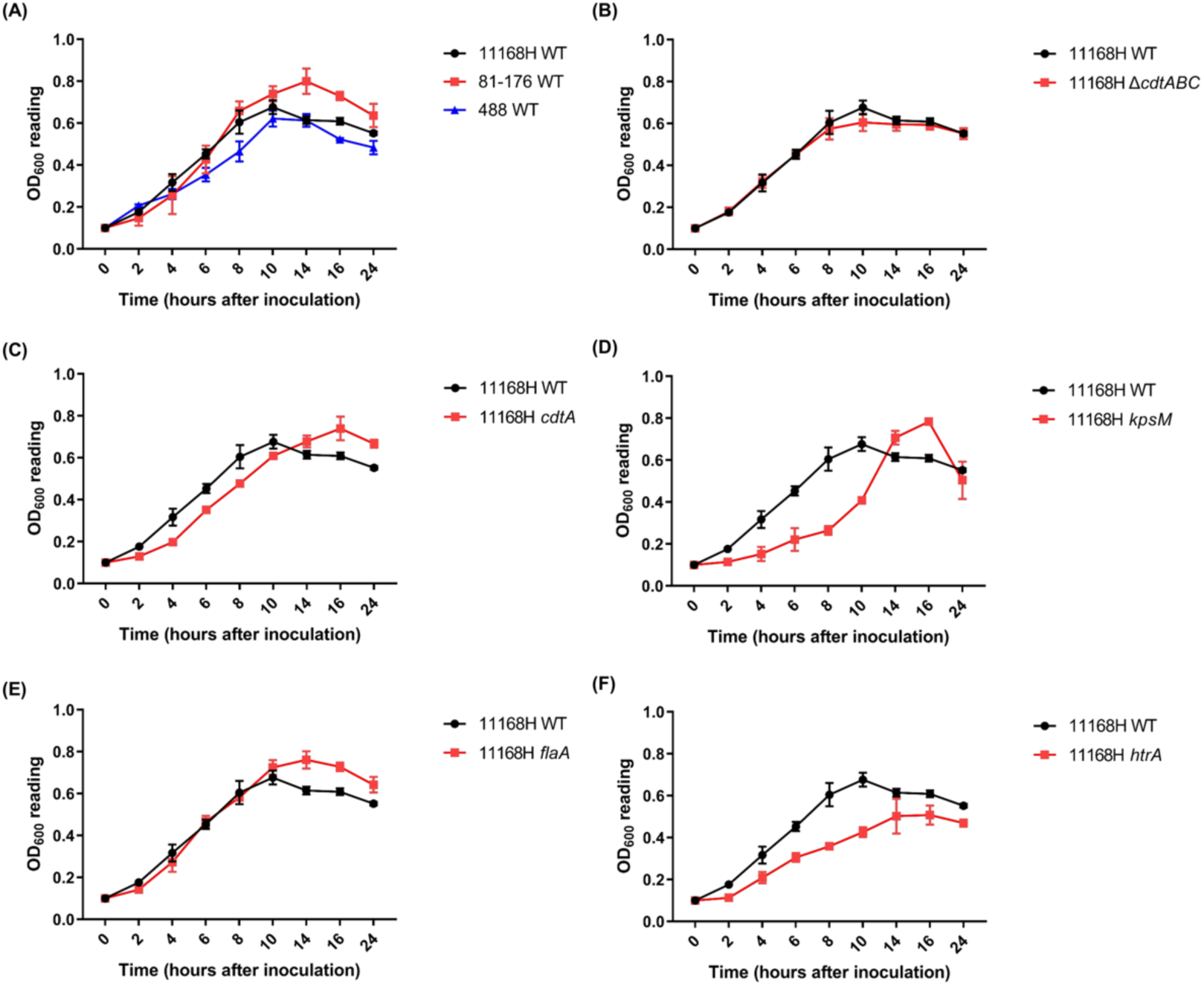
Growth kinetics of (A) *C. jejuni* 11168H, 81-176 and 488 wild-type strains (WT) and *C. jejuni* 11168H wild-type strain and (B) *cdtABC* operon, (C) *cdtA*, (D) *kpsM*, (E) *flaA* and (F) *htrA*. *C. jejuni* strains were inoculated in *Brucella* broth resulting in OD_600_ of 0.1. The inoculated flasks were incubated with shaking at 75 rpm at 37°C under microaerobic conditions. The OD_600_ of inoculated flasks was recorded at 2, 4, 6, 8, 10, 14, 16, and 24 h after inoculation.

**S6 Fig.**
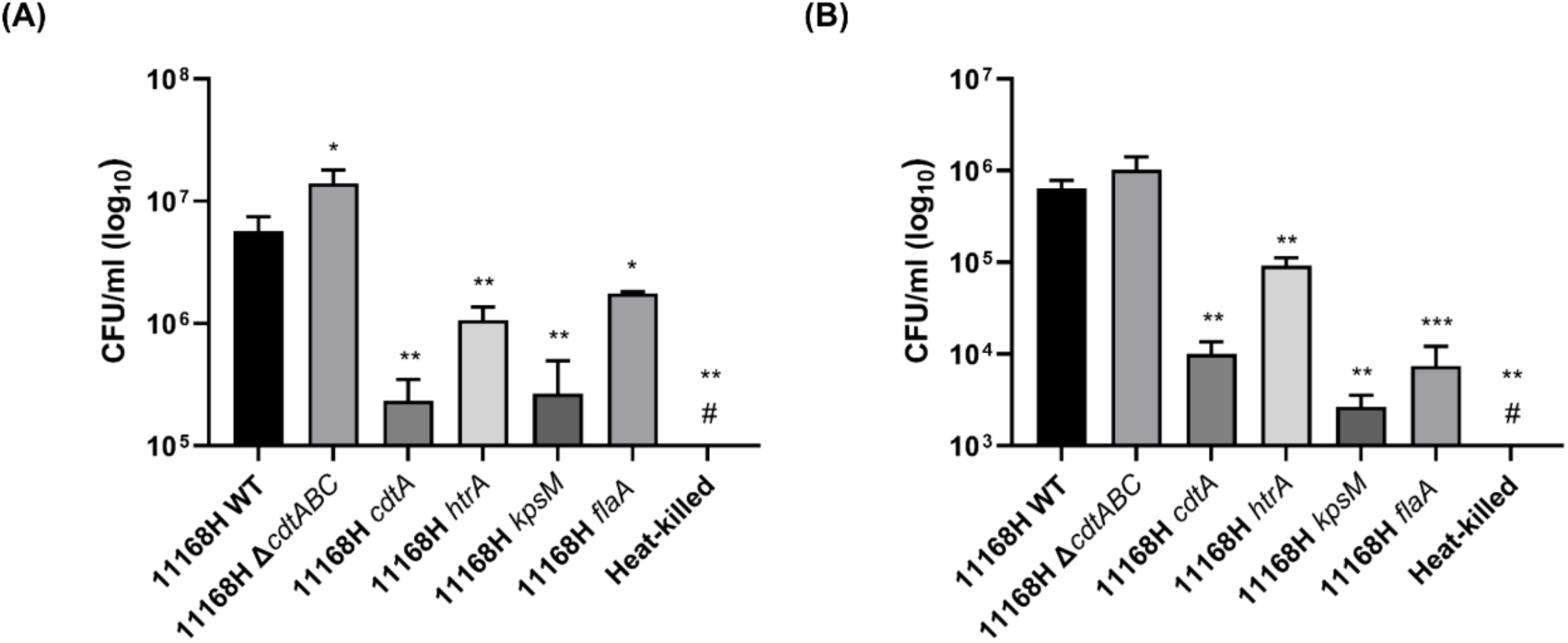
**Interactions with and invasion of T84 intestinal epithelial cells by *C. jejuni* 11168H wild-type strain and different mutants**. T84 cells were infected with *C. jejuni* 11168H wild-type strain, 11168H mutants or heat-killed 11168H wild-type strain for 3 h at 37°C in a 5% CO_2_ incubator (MOI of 200:1). (A) For interaction assays, T84 cells were then washed with PBS and lysed with 0.1% (v/v) Triton X-100 and CFU/ml were recorded after incubation. (B) For invasion assays, T84 cells were then incubated with gentamicin (150 µg/ml) for 2 h to kill extracellular bacteria and then lysed with 0.1% (v/v) Triton X-100 and CFU/ml were recorded after incubation. Three biological and three technical replicates were performed for each experiment. # denotes no growth was observed. Asterisks denote a statistically significant difference (* = *p* < 0.05; ** = *p* < 0.01; *** = *p* < 0.001).

